# Lipid-siRNA conjugate accesses perivascular transport and achieves durable knockdown throughout the central nervous system

**DOI:** 10.1101/2024.06.09.598079

**Authors:** Alexander G. Sorets, Katrina R. Schwensen, Nora Francini, Andrew Kjar, Adam M. Abdulrahman, Alena Shostak, Ketaki A. Katdare, Kathleen M. Schoch, Rebecca P. Cowell, Joshua C. Park, Alexander P. Ligocki, William T. Ford, Lissa Ventura-Antunes, Ella N. Hoogenboezem, Alex Prusky, Mark Castleberry, Danielle L. Michell, Timothy M. Miller, Kasey C. Vickers, Matthew S. Schrag, Craig L. Duvall, Ethan S. Lippmann

## Abstract

Short-interfering RNA (siRNA) has gained significant interest for treatment of neurological diseases by providing the capacity to achieve sustained inhibition of nearly any gene target. Yet, efficacious drug delivery throughout deep brain structures of the CNS remains a considerable hurdle for intrathecally administered therapeutics. We herein describe an albumin-binding lipid-siRNA conjugate that transports along meningeal and perivascular CSF pathways, leading to broad dispersion throughout the CNS parenchyma. We provide a detailed examination of the temporal kinetics of gene silencing, highlighting potent knockdown for up to five months from a single injection without detectable toxicity. Single-cell RNA sequencing further demonstrates gene silencing activity across diverse cell populations in the parenchyma and at brain borders, which may provide new avenues for neurological disease-modifying therapies.

## Introduction

Short interfering RNA (siRNA) conjugates are gaining clinical traction with the recent FDA approval of four carrier-free, trivalent GalNAc conjugates that act in the liver.^1^ There is a now a push to develop siRNA technology to treat neurodegenerative diseases, which requires effective delivery throughout central nervous system (CNS) tissues without causing prohibitive side effects. Clinical use of nucleic acid therapeutics, such as antisense oligonucleotides (ASOs) and siRNAs, is powered by advances in nucleic acid chemical modifications that enable high metabolic stability, intracellular delivery, and sustained, selective target inhibition *in vivo*.^2^ Unfortunately, at present, systemic administration of siRNA-based therapeutics do not reach the CNS in meaningful concentrations due to the blood-brain barrier (BBB).^3^ As an alternative, direct intrathecal administration can deliver therapeutics to cerebrospinal fluid (CSF) space, with the goal of achieving widespread delivery across CSF to brain interfaces.^4^ Notably, several ASO therapies targeting genes in the spinal cord provide clinical benefit after intrathecal administration; Spinraza and Tofersen have extended survival for patients with Spinal Muscular Atrophy and Amyotrophic Lateral Sclerosis, respectively.^5,6^

Even with intrathecal drug delivery, the limited interface between bulk CSF circulation and brain parenchyma poses a considerable challenge for achieving therapeutic effects beyond the local environment and the superficial layers of the brain. In mammals, CSF is secreted into the ventricles by the choroid plexus and most CSF is contained in the subarachnoid space surrounding the brain and spinal cord, separated from the parenchyma by the pia mater and glia limitans superficialis (∼20 nm gaps between adjacent astrocytic endfeet).^7^ If therapeutics do traverse these barriers to reach the parenchyma, compounds typically only diffuse into superficial brain structures, and diffusional transport alone is not efficient for delivery over the long distances needed to reach deeper brain structures such as the striatum.^8^ The unmet need for striatal delivery is highlighted by unsuccessful intrathecally-administered ASO therapies for Huntington’s disease (HD). These clinical trials were based in part on ASO studies that achieved potent huntingtin (*HTT*) mRNA silencing in the spinal cord and cortex of non-human primates, but not in the striatum.^9^ Despite the high dose administered, it is unclear if the drug was effectively delivered to the relevant deep neuroanatomical structures to engage its target, and the trial did not reach its efficacy endpoints.^10^

Leveraging convective CSF flow along perivascular spaces could overcome diffusional limitations and provide a direct, mechanism for transport to deep brain structures.^11–13^ Perivascular CSF-containing compartments sheath all large cerebral vessels extending from the subarachnoid space to deep tissue, creating an extensive network that is thought to facilitate the exchange of molecules between CSF and brain interstitial fluid.^11,14^ Intriguingly, multiple orthogonal studies suggest that the striatum receives the most perivascular CSF flow, highlighting the potential of this transport pathway for accessing deep brain structures.^15,16^ While perivascular transport has been investigated for antibody-based therapies^8^ and ASOs,^17^ it has not been examined for siRNA. Another benefit of perivascular delivery is the potential to therapeutically target cell populations such as border-associated macrophages (BAMs)^18,19^ and fibroblasts^20,21^ that reside in this compartment and are becoming increasingly implicated in neurodegeneration.

Beyond reaching the parenchyma, other delivery challenges for nucleic acid drugs include their tissue dispersion and cellular uptake. Structural design optimization for these outcomes is further exacerbated by the fact that these two processes are typically at odds with one another. For example, unconjugated siRNA structures administered carrier-free into CSF are able to disperse homogenously into the CNS but are not sufficiently internalized by cells to achieve sustained gene silencing.^22^ Conjugation of siRNA to lipophilic moieties is intended to improve cellular uptake through non-specific membrane interactions.^23,24^ However, cellular uptake can also counter transport; for example, cholesterol to an siRNA promotes robust cellular association, but generates steep concentration gradients around CSF-brain interfaces instead of achieving more homogeneous distribution within the brain.^25^ In addition to reducing penetration and delivery homogeneity, high local concentrations of cholesterol are toxic.^26,27^ Thus, to advance clinically relevant lipid-siRNA biotechnology, a balance must be achieved between transport through CSF compartments (including perivascular spaces), dispersion through parenchymal tissue, and sufficient cellular uptake, while avoiding overt toxicity.

Here, we hypothesized that a lipid-siRNA conjugate that reversibly binds to albumin would achieve an ideal combination of brain tissue penetration along with effective intracellular delivery and durable, on-target gene silencing. Prior studies have shown that albumin-binding dyes have reduced clearance to the periphery after intracerebroventricular (ICV) infusion in mice^28^ and accumulate in perivascular compartments after ICV injection in rats.^15,29^ We recently developed a lipid-siRNA conjugate termed “L2-siRNA” that was optimized in the context of intravenous delivery for albumin hitchhiking toward delivery to tumors.^30^ This work studied the effects of linker and lipid structure on the ability of the siRNA to navigate both systemic- and cell-level delivery barriers. Structural features were identified that reduce conjugate micellization while tuning non-covalent albumin affinity that enables harnessing albumin as an endogenous carrier protein, while also allowing release from albumin such that the lipids can participate in cellular entry. Here, we characterized this platform for delivery into the CNS following administration into CSF, using ICV injection in mice and intrathecal injection in rats. We investigated L2-siRNA access to perivascular transport pathways, spatial variation in brain delivery, and target gene silencing potency and longevity. Single cell-RNA sequencing (scRNA-seq) was applied to examine cell type dependent conjugate gene silencing activity. Lastly, building on the emerging role of CSF-interfacing brain border cells in health and disease, we completed a unique and deep exploration of the ability of L2-siRNA to silence genes in leptomeningeal fibroblasts, choroid plexus epithelial cells, and BAMs. Collectively, this work establishes L2-siRNA as a promising platform for gene silencing in the CNS, while also rigorously characterizing its potential for targeting specific cell types and regions of the brain.

## Results

### Lipid-siRNA conjugate structure and properties

To initially examine how the properties of L2-siRNA might impact CSF to brain delivery, we evaluated *in vitro* uptake in neuroblastoma cells and *ex vivo* association with albumin in CSF, with comparisons to unconjugated siRNA (“siRNA”) and cholesterol-conjugated siRNA (“Chol-siRNA”). The blunt-ended siRNA design contains alternating 2ʹOMe and 2ʹF on both strands in a “zipper” pattern. Phosphorothioate (PS) linkages were used between the last two bases on the ends of both strands, and vinyl phosphonate was integrated at the 5ʹ antisense position, which is an important feature for maximal silencing in the CNS.^22,31^ Structurally, the L2-siRNA branches off the 5’ end of the sense strand and contains a spacer (18-ethylene glycol repeats) followed by divalent lipid tails (two 18-carbon stearyl groups) (Fig. 1A). More L2-siRNA was taken up than the parent siRNA, while Chol-siRNA, a benchmark chosen for its known cell penetration capacity,^24^ had the highest level of uptake in serum-free media (Supplementary Fig. 1A). Despite uptake differences, L2-siRNA carrier-free gene silencing activity was equivalent to Chol-siRNA at 250 nM (Supplementary Fig. 1B). In addition, we confirmed L2-siRNA association with albumin in *ex vivo* human CSF, as demonstrated by fast protein liquid chromatography (FPLC). In comparison, Chol-siRNA and unconjugated siRNA showed no association with albumin (Supplementary Fig. 1C,D). These outcomes suggested that L2-siRNA is a good candidate to potentially leverage albumin for perivascular transport, while also having sufficient capacity for intracellular delivery within the CNS.

**Figure 1:**
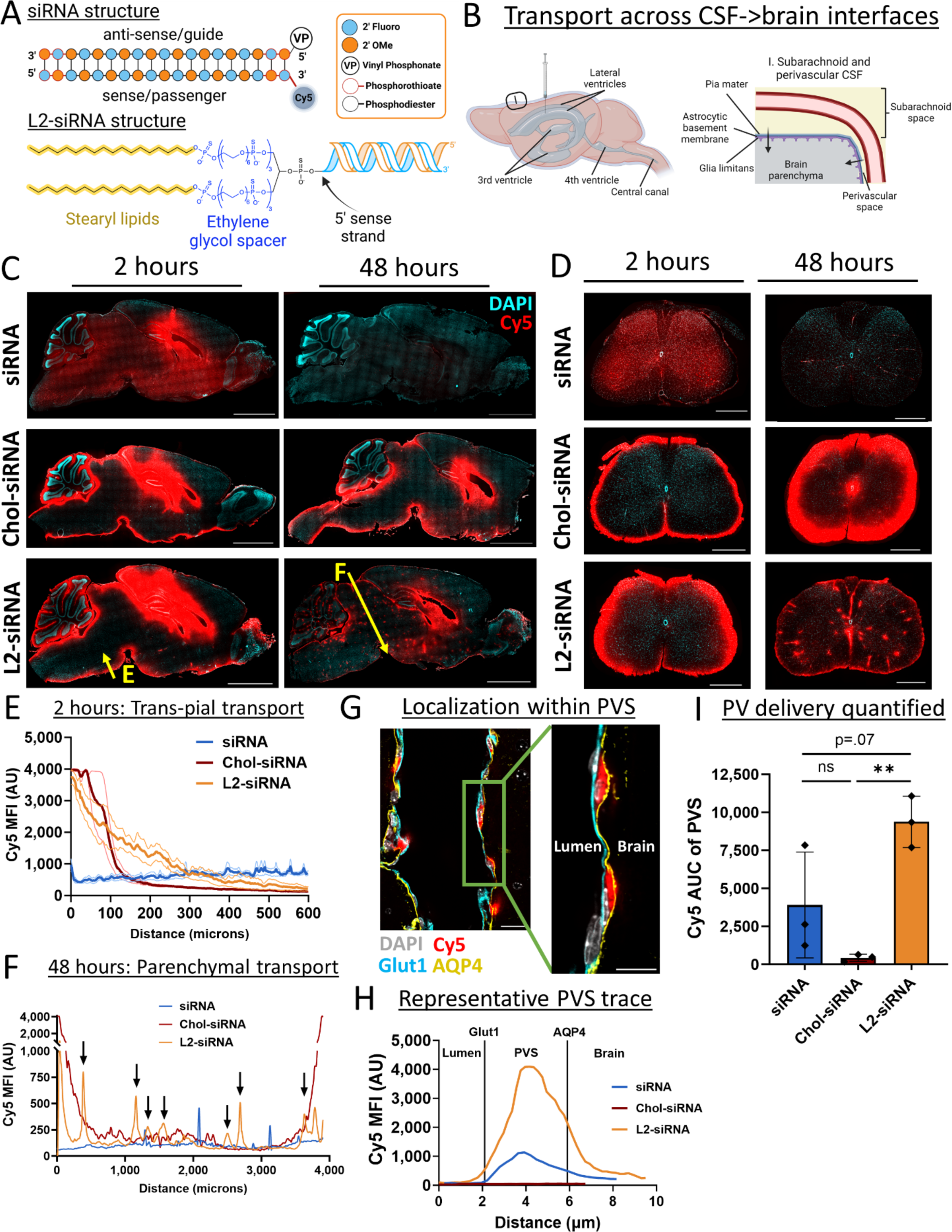
Lipid-siRNA conjugate structure determines CSF to brain transport. A. All nucleic acid structures are synthesized using standard zipper siRNA chemistry, characterized by alternating 2′ Omethyl (OMe) and fluoro bases, terminating with two phosphorothioate bonds. All sequences utilize the 5’ vinyl phosphonate (VP) on the antisense strand, and when applicable the Cy5 fluorophore is positioned at the 3’ position of the sense strand. Lipid conjugates are attached to the 5’ end of the sense strand. L2-siRNA is composed of a divalent spacer and lipid tail. The spacer contains three repeats of hexaethylene glycol groups on each side of a brancher (totaling 18 ethylene glycol units), and the lipid is an 18-carbon alkyl chain. Chol-siRNA (structure not shown) contains a triethylene glycol spacer separating the cholesterol moiety from the siRNA. B. Overview of CSF anatomy highlighting interfaces for drug delivery. To examine biodistribution, Cy5-tagged compounds are injected into CSF through the intracerebroventricular route. CSF flows from the ventricular system to the subarachnoid space, and then some CSF moves alongside vessels in the perivascular spaces of the brain. C. Histological examination of CNS distribution either 2 or 48 hours after ICV injection of unconjugated siRNA, Chol-siRNA, or L2-siRNA (7.5-10 nmol dose). Cy5 signal is presented in red and DAPI nuclear signal is presented in cyan. Representative sagittal images of left hemispheres are displayed (N=3 mice per condition). Section thickness = 30 μm, scale bars = 2.5 mm. D. Histological examination of distribution in spinal cord either 2 or 48 hours after ICV injection of unconjugated siRNA, Chol-siRNA, or L2-siRNA (7.5-10 nmol dose). The 2-hour time point shows representative lumbar regions while the 48-hour time point shows representative cervical regions (N=3 mice per condition). Section thickness = 30 μm, scale bars = 500 µm. E. Plot profile of Cy5 signal around the ventral subarachnoid CSF 2 hours after injection. Yellow arrow in panel C (labeled as “E”) represents the analyzed region. Data are presented from N=3 biological replicates (faint lines are ± SEM and solid line is the mean). F. Representative plot profile of Cy5 signal across deeper brain structures 48 hours after injection (from midbrain to interpeduncular cistern). Yellow arrow in panel C (labeled as “F”) represents the analyzed region. Arrows point to L2-siRNA peaks reflecting perivascular transport far from brain boundaries. Additional biological replicates are shown in Supplementary Figure 3. G. Representative image showing perivascular localization of L2-siRNA 48 hours after injection. Perivascular space is defined as the region between Glut1+ endothelium and AQP4+ glia limitans. Scale bar = 20 µm (left) or 10 µm (right). The inset is acquired at the indicated area but at a slightly different z-plane using confocal microscopy. H. Representative traces of Cy5 fluorescence within the perivascular space (PVS) shown for a representative vessel from each condition, with boundaries demarcated by peak Glut1 and Aqp4 intensities. I. Quantification of perivascular Cy5 signal. Each dot represents the average of 5 non-capillary vessels from a single mouse. N=3 biological replicates per condition and data presented as mean ± SD. Statistical significance was calculated using a one-way ANOVA with Bonferroni’s correction (** p<0.01, ns – not significant). AU = arbitrary units, AUC = area under curve.

### L2-siRNA traffics in perivascular spaces and enters the brain by diffusion from the subarachnoid compartment

Considering the brain anatomy (Fig. 1B), we first sought to investigate how lipid properties of siRNA conjugates impact CSF to brain transport *in vivo*. We injected mice ICV with Cy5-labeled unconjugated siRNA, Chol-siRNA, or L2-siRNA and examined biodistribution after 2 and 48 hours. We observed that unconjugated siRNA rapidly disperses through the mouse brain and spinal cord by 2 hours post-administration, but that the siRNA was minimally retained in the CNS by 48-hours (Fig. 1C,D). Coupled with the observation of accumulation of siRNA in kidneys at 2 hours (Supplementary Fig. 2), these results suggest that unconjugated siRNA is not efficiently internalized by CNS cells causing it to be eliminated quickly from the CNS. In contrast, Chol-siRNA exhibited high brain retention at both timepoints, but it was characterized by steep concentration gradients away from the CSF-brain interface (Fig. 1C). This delivery pattern was also present in the spinal cord, where Chol-siRNA exhibited steep gradients from the subarachnoid space toward the central canal (Fig. 1D). In contrast, L2-siRNA appeared within all CSF interfaces in the parenchyma (Fig. 1C): ventricular (trans-ependymal), subarachnoid (trans-pial, including cisternal; Fig. 1E), and perivascular. In the spinal cord, L2-siRNA also exhibited broad tissue dispersion along with observable puncta (Fig. 1D), apparently associated with perivascular compartments. To demonstrate reproducibility, an additional experimental replicate for 48-hour delivery in brain is shown in Supplementary Fig. 3.

We next more definitively assessed the localization of L2-siRNA in perivascular spaces. First, we measured the Cy5 intensity profile across the midbrain to the interpeduncular cistern (far from injection site) and detected peaks of L2-siRNA consistent with perivascular transport to deep brain structures (Fig. 1F). Localization within the perivascular space was confirmed by the presence of L2-siRNA between endothelial cells (Glut1+) and the astrocytic parenchymal glia limitans (Aquaporin-4+) (Fig. 1G).

Measuring Cy5 intensity between these markers confirmed that L2-siRNA achieves greater perivascular delivery than Chol-siRNA (Fig. 1H,I). To evaluate whether delivery to perivascular spaces could be an artifact of the injection volume (e.g. pressure-driven effects), we injected mice ICV with either 2 μl or 10 μl of L2-siRNA at equivalent total dose (2 nmol). We found that L2-siRNA similarly accessed perivascular spaces in the brain and spinal cord when injected in the lower volume (Supplementary Fig. 4), suggesting the molecular properties of L2-siRNA are responsible for observed outcomes. These results indicate that L2-siRNA delivered into the CSF is trafficked through perivascular spaces, providing a route for transport into deeper parenchymal structures.

### L2-siRNA achieves durable gene and protein silencing in bulk tissue across multiple brain regions

Based on the desirable (dispersed yet retained) delivery profile of L2-siRNA, we next sought to characterize potency and kinetics of target gene silencing. We chose *Htt* as a disease-relevant target that is ubiquitously expressed by CNS cells and for which a well-validated siRNA sequence exists (“Htt10150”).^25^ Targeting *Htt* also permits indirect comparisons to previous studies. Adult mice were injected with 15 nmol via ICV route. This is a relatively low dose compared to other nucleic acid therapeutics targeting *Htt* in mice, such as a divalent siRNA structure (40 nmol)^31^ or clinical ASO (∼ 100 nmol administered over two weeks).^9^ We measured *Htt* mRNA by RT-qPCR, HTT protein by Western blotting, and absolute amount of antisense strand delivery with a peptide nucleic acid (PNA) hybridization assay; analyses were performed for CNS regions both proximal to the lateral ventricle injection site (striatum, hippocampus, cortex) as well as distal to the injection (brainstem, cerebellum) (Fig. 2A). To identify a suitable control, we assessed *Htt* expression in these brain regions after ICV delivery of vehicle (0.9% saline) or a non-targeting siRNA conjugated to the L2 lipid (termed L2-siRNA^NTC^, targeting luciferase, which is not expressed in mice). We determined that treatment with L2-siRNA^NTC^ did not change *Htt* expression appreciably compared to vehicle (Supplementary Fig. 5) and therefore proceeded with L2-siRNA^NTC^ as the primary negative control for gene silencing studies.

**Figure 2:**
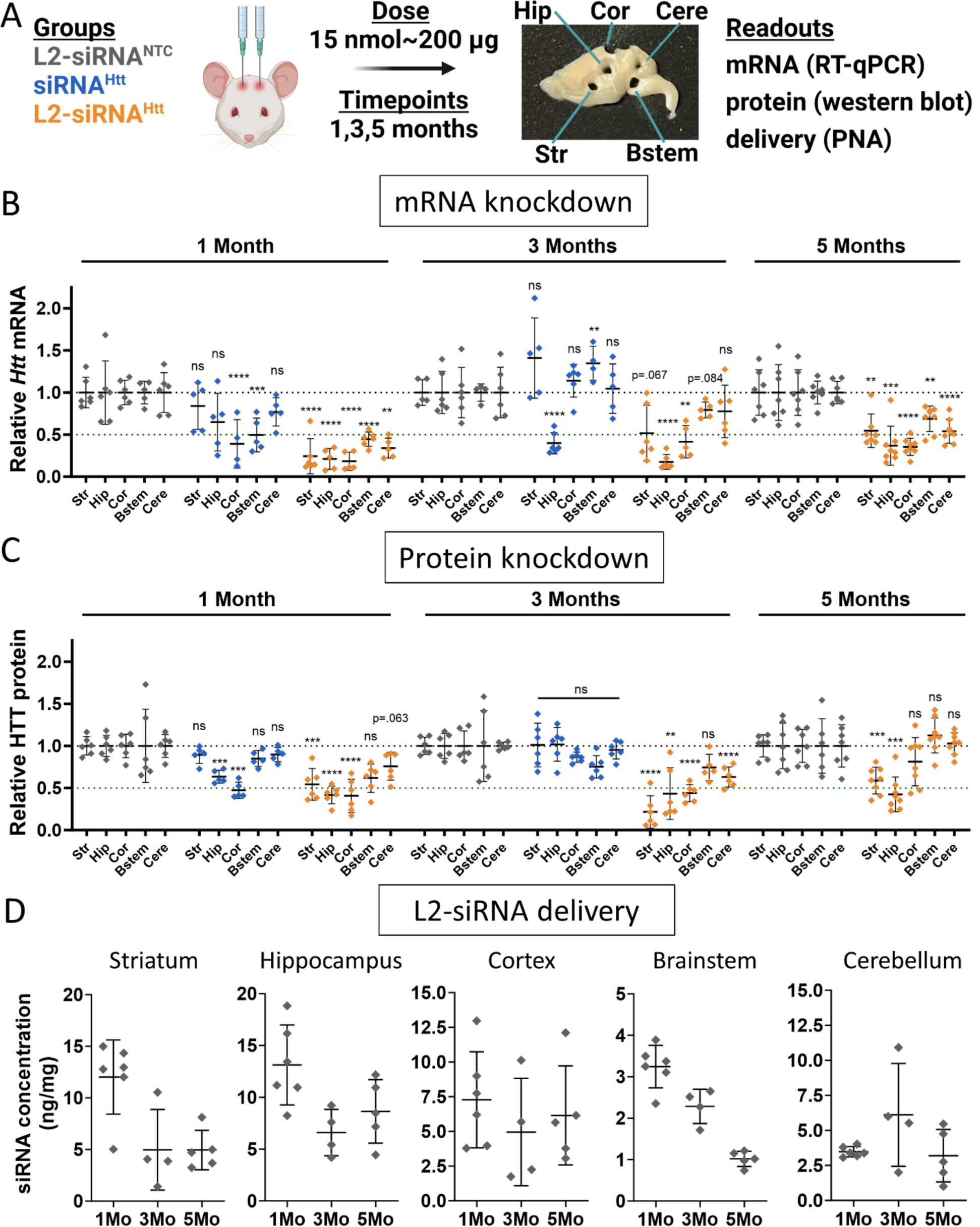
L2-siRNA^Htt^ exhibits potent and sustained gene and protein silencing throughout the parenchyma. A. Experimental approach for evaluating gene silencing. Mice receive a bilateral ICV injection of 15 nmol L2-siRNA^NTC^, siRNA^Htt^, or L2-siRNA^Htt^ and brain tissue is harvested after 1, 3, or 5 months. The brain is sliced into 1 mm sagittal sections, which are biopsy punched (2 mm) to extract different regions for analysis. A representative brain section is shown with biopsy punches extracted. RT-qPCR samples are acquired from the first 1 mm medial slice of the left hemisphere, western blot samples are acquired from the second medial slice of the left hemisphere, and PNA samples are acquired from the first medial slice of the right hemisphere. Str, striatum; Hip, hippocampus; Cor, cortex; Bstem, brainstem; Cere, cerebellum. B. *Htt* mRNA levels in parenchymal regions as measured by RT-qPCR. Each region and time point is normalized to its respective L2-siRNA^NTC^ control. For 1 and 3 months, significance was calculated for each region and timepoint as a one-way ANOVA compared to L2-siRNA^NTC^ with Bonferroni’s correction for multiple comparisons (N=5-6 mice). For 5 months, unpaired two-tailed t-tests were performed for each region (N=7-8 mice). C. HTT protein levels in parenchymal regions as measured by western blot. Each region and time point is normalized to its respective L2-siRNA^NTC^ control. Raw western blots are shown in Supplementary Figure 6. For 1 and 3 months, significance was calculated for each region and timepoint as a one-way ANOVA compared to L2-siRNA^NTC^ with Bonferroni’s correction for multiple comparisons (N=5-6 mice). For 5 months, an unpaired two-tailed t-test was performed for each region (N=7-8 mice). D. Absolute amount of antisense strand (ng) per milligram of brain tissue as measured by the PNA assay for L2-siRNA^NTC^. Detailed description of quantification is demonstrated in Supplementary Figure 7. N=4-6 biological replicates per region per time point (M=month). For all panels, data are represented as mean ± SD and each point represents an individual biological replicate (i.e., a single mouse brain). Mo = months. In all panels, statistical significance has the same labels (* p<0.05, ** p<0.01, *** p<0.001, **** p<0.0001, ns – not significant).

At 1 month after injection, *Htt*-targeting L2-siRNA (referred to as L2-siRNA^Htt^) demonstrated potent mRNA knockdown (>50%) in all brain regions tested, whereas unconjugated siRNA (referred to as siRNA^Htt^) only exhibited silencing in the cortex and brainstem, highlighting the benefit of L2 conjugation (Fig. 2B). Similarly, at the protein level, siRNA^Htt^ mediated some silencing in the cortex and hippocampus, but less than L2-siRNA^Htt^, which also induced robust knockdown in the striatum (Fig. 2C and Supplementary Fig. 6). At 3 months post-injection, mice receiving siRNA^Htt^ no longer exhibited significant protein knockdown in any regions, and only the hippocampus exhibited a reduction in the target mRNA. In contrast, mice receiving L2-siRNA^Htt^ exhibited sustained gene and protein knockdown; we note significant reduction in HTT protein at 3 months in the striatum, hippocampus, cortex, and cerebellum. Impressively, at a prolonged 5-month timepoint, L2-siRNA^Htt^ still achieved mRNA silencing in all examined parenchymal regions. The protein levels were also reduced by L2-siRNA^Htt^ treatment in some regions (striatum, hippocampus), while others returned to basal levels. Unconjugated siRNA^Htt^ was omitted from the 5-month study since its activity was lost at 3 months. Overall, our data show that L2-siRNA^Htt^ potentiates widespread and prolonged gene and protein silencing throughout the CNS compared to the parent siRNA.

To define the relationship between siRNA delivery and gene target knockdown, we quantified L2-siRNA present over a time course using a PNA hybridization assay to measure mass of antisense strand per tissue mass (Fig. 2D and Supplementary Fig. 7). Consistently, regions with the highest delivery (striatum, hippocampus, and cortex) also had the most gene silencing at all timepoints. While siRNA accumulation decreased over time (as expected), a considerable amount of siRNA-L2 was still present at 3 and 5 months, consistent with the recently-posited endolysosomal depot phenomenon.^32^ This evidence suggests that, once delivered, this siRNA complex remains active in tissue for a prolonged period, a feature that is advantageous for minimizing treatment frequency in chronic diseases.

We also compared L2-siRNA^Htt^ to a *Htt*-targeting ASO. To match our experimental regime, we administered an equimolar ICV dose (15 nmol, ∼ 95 μg) of the “MoHu” gapmer ASO^Htt^ that was designed with homology for mouse and human *HTT*.^9^ Gene silencing was assessed at a 1-month time point, and given the disparate structures between the two molecules, normalization was performed relative to a vehicle control. L2-siRNA^Htt^ was more potent than ASO^Htt^ administered equimolar dose, demonstrating statistically significant enhancement of mRNA knockdown in all parenchyma and spinal cord regions except hippocampus, where both compounds caused robust knockdown (Supplementary Fig. 8). Of particular note, L2-siRNA^Htt^ was more effective in the striatum (∼80% reduction) than ASO^Htt^ (∼25% reduction), which is highly relevant considering the striatum is the region predominantly affected by Huntington’s disease.

### L2-siRNA achieves durable gene and protein silencing throughout the spinal cord

The spinal cord is directly connected to the CSF circulation and implicated in several diseases targetable by nucleic acid therapies. However, unlike the intrathecal route of administration often used to modulate disease targets of the spinal cord, delivery to the spinal cord requires greater transport from an ICV injection site. Evaluation of the cervical, thoracic, and lumbar spinal cord regions at 1-, 3-, and 5-months post-injection by the PNA assay showed that L2-siRNA levels in the spinal cord decrease linearly over time from ∼3 ng/mg to ∼1 ng/mg (Fig. 3C,D). These levels are lower than the parenchyma, however, L2-siRNA^Htt^ gene silencing activity remained high in all segments of the spinal cord at both the mRNA and protein levels, whereas unconjugated siRNA did not mediate knockdown of HTT protein (Fig. 3A,B). These outcomes further highlight the effectiveness of L2-siRNA throughout the CNS.

**Figure 3:**
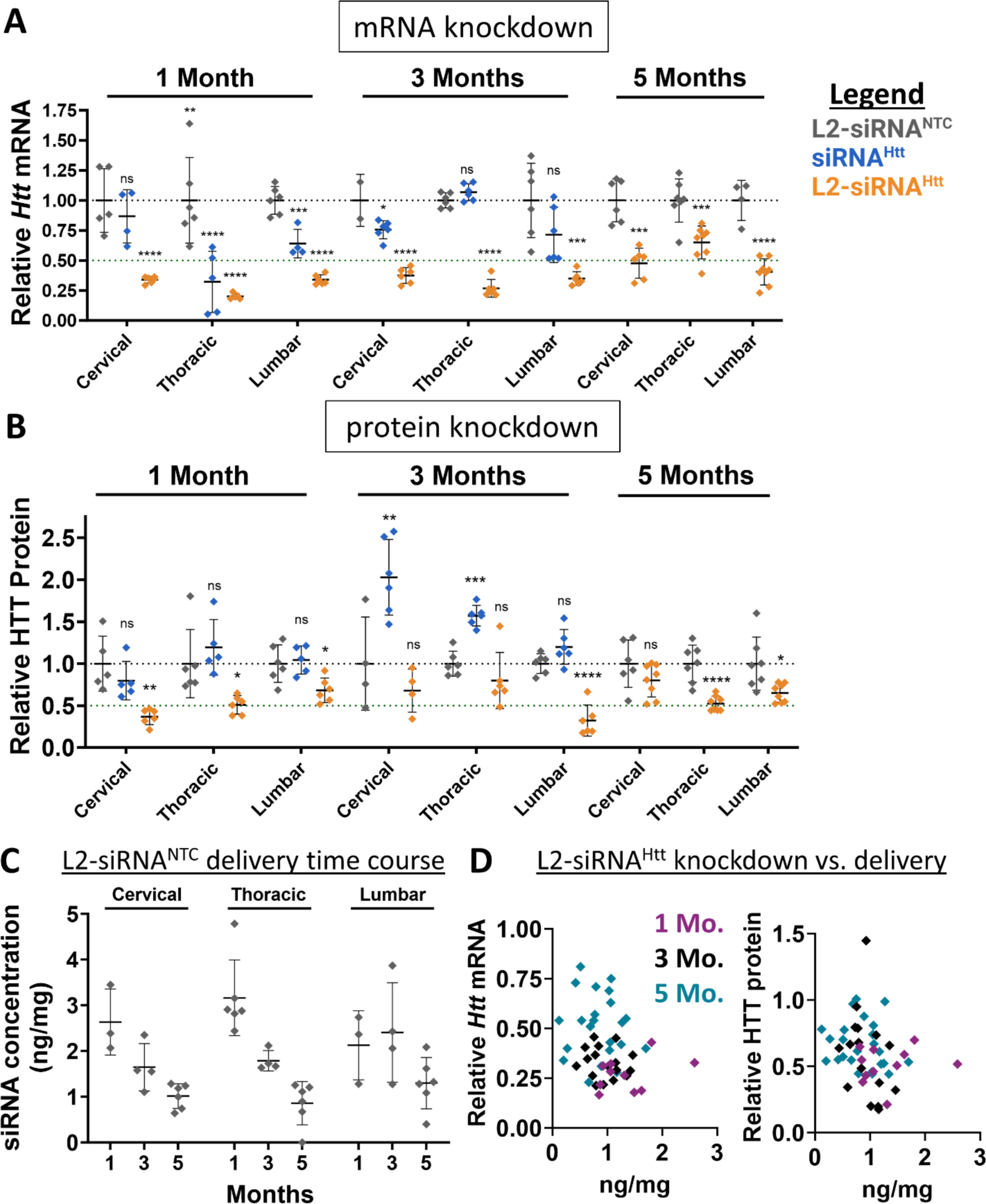
L2-siRNA exhibits potent and sustained mRNA and protein silencing in the spinal cord. A. *Htt* expression levels in spinal cord regions as measured by RT-qPCR. Mice receive a bilateral ICV injection (15 nmol total dose) of L2-siRNA^NTC^, siRNA^Htt^, or L2-siRNA^Htt^ and the spinal cords were collected after 1, 3, or 5 months. N=4-6 spinal cords analyzed at 1 and 3 months, and N=7-8 spinal cords analyzed at 5 months. For 1 and 3 months, significance was calculated for each region and timepoint as a one-way ANOVA compared to the L2-siRNA^NTC^ control with Bonferroni’s correction for multiple comparisons. For 5 months, an unpaired two-tailed t-test was performed for each region. B. HTT expression levels in spinal cord regions as measured by western blot. N=4-6 spinal cords analyzed at 1 and 3 months, and N=7-8 spinal cord analyzed at 5 months. For 1 and 3 months, significance was calculated for each region and timepoint as a one-way ANOVA compared to the L2-siRNA^NTC^ control with Bonferroni’s correction for multiple comparisons. For 5 months, unpaired two-tailed t-test was performed for each region. C. Absolute amount of antisense strand (ng) per milligram of spinal cord tissue measured by the PNA assay at 1, 3, and 5 months (M). N=3-6 biological replicates per region per time point. Values below the limit of detection were plotted as 0. D. Relationship between L2-siRNA delivery and gene/protein knockdown. For every graph, each point represents an individual biological replicate (i.e., a single mouse spinal cord) and data are presented as mean ± SD. Mo. = months. In all panels, statistical significance has the same labels (* p<0.05, ** p<0.01, *** p<0.001, **** p<0.0001, ns – not significant).

### L2-siRNA effectively targets diverse CNS cell types

Next, cell type-specific uptake and gene silencing activity were characterized for L2-siRNA. To assess cellular uptake, mice were administered Cy5-tagged L2-siRNA or parent siRNA (10 nmol) by ICV, and flow cytometry was performed after 48 hours to measure Cy5 levels in various cell types defined as: Thy1+ neurons, ACSA2+ astrocytes, CD11b+ microglia/macrophages, and CD31+ endothelial cells. Oligodendrocytes were removed from analysis based on O1+ staining, as myelin can promote non-specific binding.^33^ We found that total and cell-specific uptake were increased by L2 conjugation (Supplementary Fig. 9), in agreement with our prior *in vitro* data (Supplementary Fig. 1A). L2-siRNA uptake into neurons was the lowest out of the cells examined, but this may have been due to poor neuron recovery in our dissociation protocol (<2% of total population). Microglia, which are resident CNS phagocytes, showed the highest L2-siRNA uptake (Supplementary Fig. 9).

To determine whether L2-siRNA uptake corresponds to functional gene silencing across CNS populations, we evaluated mRNA knockdown using scRNA-seq. Here, we used an siRNA sequence targeting *Ppib* because this gene is abundantly and ubiquitously expressed across CNS cell types.^34^ Potent siRNA sequences have also been designed and validated against *Ppib* elsewhere.^35^ Matching the experimental parameters of bulk tissue knockdown analyses, we injected 15 nmol of either L2-siRNA*^Ppib^* or L2-siRNA^NTC^ and collected cells after 1 month for evaluation. Since we observed substantial uptake into Cd11b+ cells by flow cytometry, we elected to enrich for myeloid cells with bead-based sorting prior to barcoding and analysis, yielding two datasets: Cd11b^pos^ cells encompassing microglia and macrophages, and Cd11b^neg^ cells comprising an untargeted sampling of all other isolated meningeal and parenchymal populations (Fig. 4A). To verify gene silencing with orthogonal methodology, cells not utilized for scRNA-seq were analyzed in parallel using qRT-PCR, which confirmed gene silencing in both Cd11b^pos^ and Cd11b^neg^ populations with an average of 45% and 65% *Ppib* knockdown, respectively (Fig. 4B). We compared RT-qPCR bulk silencing to *Ppib* knockdown measured by scRNA-seq and these outputs closely matched for both CD11b^pos^ and CD11b^neg^ populations (Fig. 4C), providing confidence in the accuracy of examining cell-specific gene silencing with scRNA-seq.

**Figure 4:**
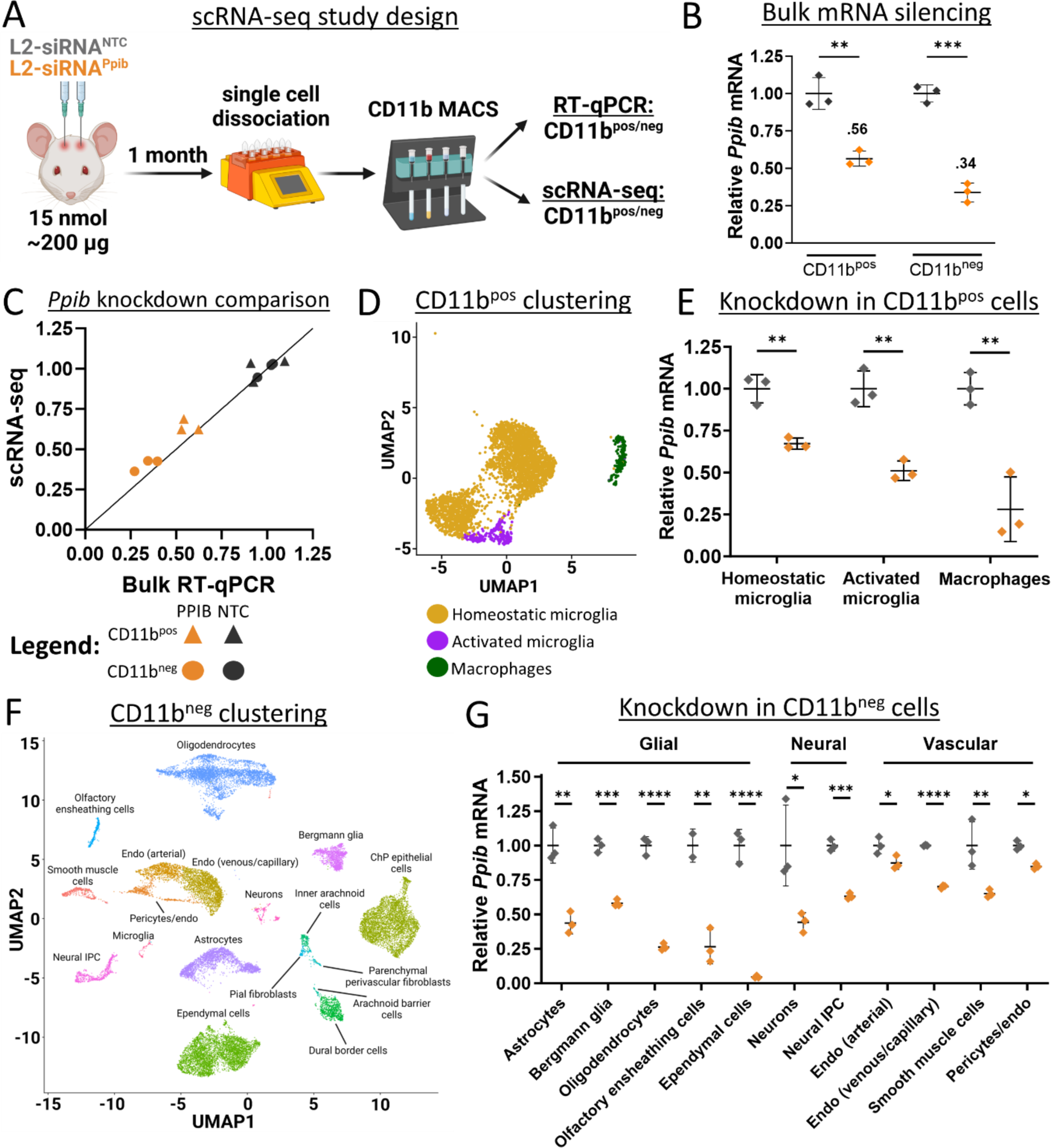
L2-siRNA potentiates gene silencing across diverse parenchymal cells. A. Experimental design for assessment of cell-specific gene silencing. Mice were administered 15 nmol of L2-siRNA^Ppib^ or L2-siRNA^NTC^, and after 1 month, brains were dissociated into single cells and bead-sorted into Cd11b^pos^ and Cd1lb^neg^ populations, which were further separated for scRNA-seq or RT-qPCR. For all experiments in this figure, N=3 individual mice (biological replicates) for both L2-siRNA^Ppib^ or L2-siRNA^NTC^. B. *Ppib* expression in Cd11b^pos^ and Cd1lb^neg^ populations measured by RT-qPCR. Statistical significance was calculated using an unpaired two-tailed t-test (** p<0.01, *** p<0.001). C. Comparison of *Ppib* expression between bulk RT-qPCR (x-axis) and scRNA-seq averaged across all identified cells (y-axis). Line with a slope of 1 represents equivalent expression between readouts. D. UMAP plot showing clustering and annotation of Cd11b^pos^ populations. E. Statistical significance was calculated using an unpaired two-tailed t-test (** p<0.01). F. UMAP dimension reduction plot showing clustering and annotation of Cd11b^neg^ populations. Unless indicated with a line, all text labels lie above their respective cluster. G. Average *Ppib* expression across different Cd11b^neg^ cell types, normalized to L2-siRNA^NTC^. Statistical significance was calculated using an unpaired two-tailed t-test (* p<0.05, ** p<0.01, *** p<0.001, **** p<0.0001, ns – not significant). Data are presented as mean ± SD in every panel.

To determine gene silencing in myeloid cells, Cd11b^pos^ cells were annotated and clustered into three populations using standard markers (Fig. 4D). Homeostatic microglia are characterized by expression of *Crybb1*, *Cst*3, *P2ry12*, *Pros1*, and *Tmem119*, while activated microglia additionally express *Apoe*, *Spp1*, *Lpl*, and *Lyz2* (Supplementary Fig. 10A,B).^36,37^ CNS macrophages include perivascular, meningeal, and choroid plexus macrophage populations, which are enriched in *Pf4*, *Lyz2*, *Ms4a7*, *Ccl24*, and *F13a1*; this population constituted ∼6% of Cd11b^pos^ cells (Supplementary Fig. 10C). Compared to the non-targeting control, L2-siRNA*^Ppib^* potentiated gene silencing in homeostatic (∼40%) and activated microglia (∼50%), as well as resident CNS macrophages (∼70%) (Fig. 4E and Supplementary Fig. 10D). Microglia are notoriously refractory to gene silencing, highlighting the significance of this result.^38^ In addition to potent myeloid gene knockdown, L2-siRNA*^Ppib^* potentiated gene silencing in a variety of other cell types identified in the Cd11b^neg^ population (Fig. 4F and Supplementary Fig. 11). For example, glial cells such as astrocytes and oligodendrocytes exhibit considerable *Ppib* knockdown, as well as ependymal cells, which line the ventricles and directly interface with CSF (Fig. 4G). Neurons exhibit modest gene silencing, and due to the low capture of this population, we could not subcluster knockdown into neuron subtypes. Endothelial cells were subclustered into venous/capillary and arterial populations (Fig. 4F); there was low but significant knockdown in arterial endothelial cells, and slightly higher silencing in the venous/capillary cells (Fig. 4G). Modest gene silencing activity in these cells is consistent with low intensity of uptake observed by flow cytometry. Overall, L2-siRNA*^Ppib^* exhibits broad gene silencing activity across diverse CNS cell types, suggesting that it can be leveraged as a platform technology for modulation of neurological diseases.

### L2-siRNA effectively targets brain border cells

Recent advances in single-cell biology have revealed the diversity of cells present at brain borders, including the choroid plexus and meninges, as well as the previously discussed perivascular spaces. There is a growing appreciation for the role brain borders play in disease,^39^ but no prior studies have rigorously evaluated siRNA delivery to the cells residing in these borders. Since brain borders directly interface with CSF flow, we investigated L2-siRNA-mediated gene silencing in these compartments. First, we examined gene silencing in leptomeningeal cells after ICV delivery. Recent characterization of fibroblast-like cells in the meninges identified five transcriptionally and spatially unique cell populations.^40^ Anatomically, the arachnoid membrane separates the dura from subarachnoid CSF and from top to bottom is composed of dural border cells, arachnoid barrier cells, and inner arachnoid cells. Beneath this membrane are pial fibroblasts, either associated with vessels (perivascular) or sitting on the pial membrane (Fig. 5A). We re-clustered our scRNA-seq data set to identify the aforementioned cells, annotating all fibroblasts outside the meninges as “parenchymal perivascular fibroblasts”, which includes perivascular, choroid plexus, and potentially median eminence fibroblasts (Fig. 5B). Impressively, knockdown was observed in all fibroblast-like populations, with dural border cells experiencing the least gene silencing, possibly because of their location on the opposing side of the arachnoid membrane. The greatest fibroblast knockdown potency was observed in cells directly contacting CSF, such as arachnoid barrier cells, pial fibroblasts, and arachnoid barrier cells (Fig. 5C).

**Figure 5:**
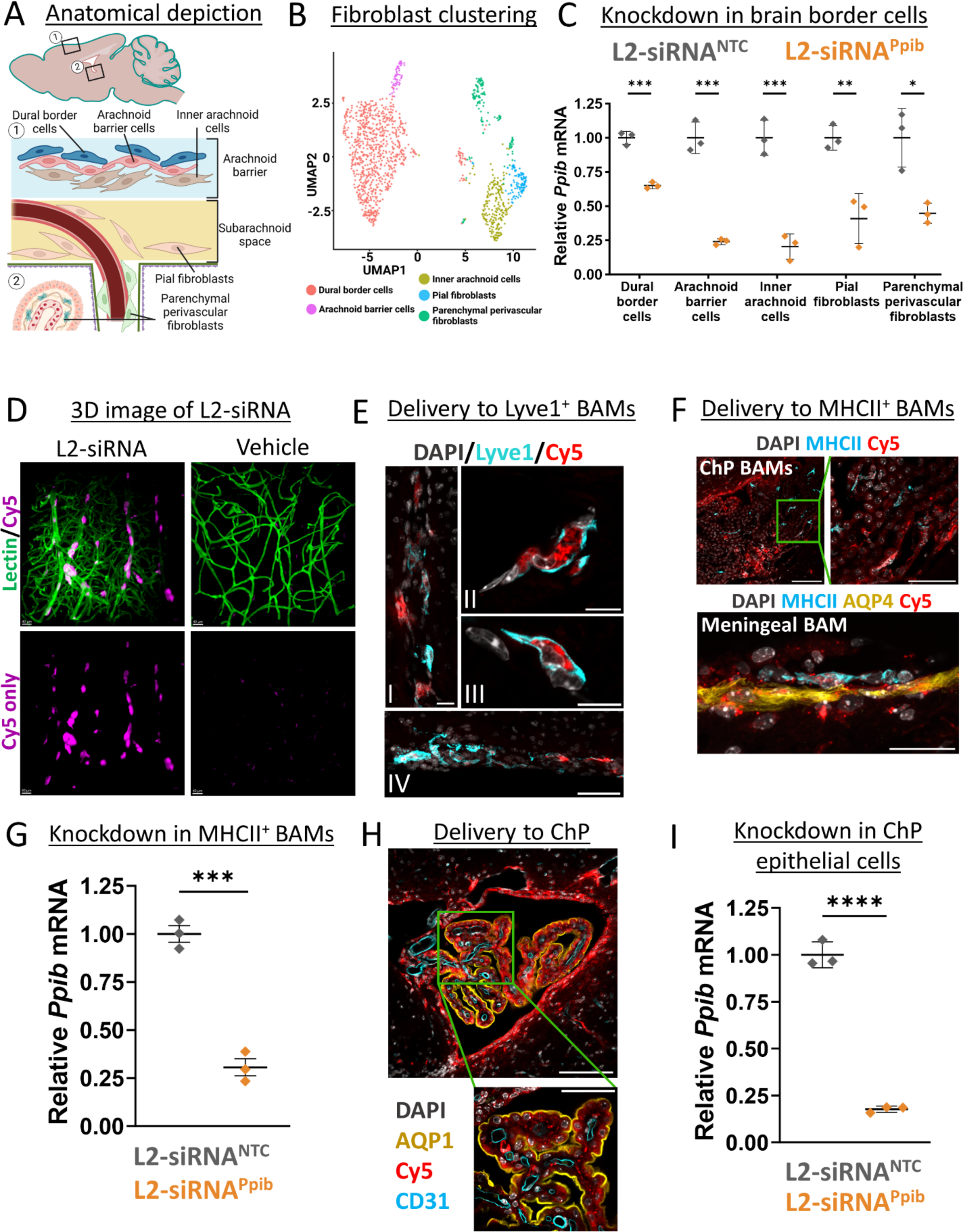
L2-siRNA exhibits effective targeting and gene silencing in brain border cells. A. Depiction of fibroblast anatomical locations in the leptomeninges. B. UMAP dimension reduction plot showing sub-clustering of fibroblast populations. C. scRNA-seq assessment of *Ppib* knockdown in fibroblasts of the brain borders at a 1-month time point. Knockdown is normalized to L2-siRNA^NTC^. N=3 individual mice (biological replicates) for both L2-siRNA*^Ppib^* and L2-siRNA^NTC^. Statistical significance was calculated using an unpaired two-tailed t-test (* p<0.05, ** p<0.01, *** p<0.001). D. Three-dimensional image demonstrating extensive perivascular delivery of L2-siRNA (magenta) around vessels (green) in the cortical surface (48 hours, 10 nmol dose). Representative images displayed are consistent across replicates. E. Representative image of Cy5-tagged L2-siRNA delivery to Lyve1^+^ BAMs 48 hours after injection. (I) Delivery along penetrating vessel, scale bar = 20 μm. (II, III) Subcellular localization in perivascular Lyve1+ BAMs, scale bar = 10 μm. (IV) Delivery to ventral meningeal Lyve1+ BAMs, scale bar = 50 μm. F. Representative images of Cy5-tagged L2-siRNA delivery to MHCII+ BAMs 48 hours after injection. Scale bar = 100 μm (top left), 50 μm (top right), 25 μm (bottom). G. scRNA-seq assessment of *Ppib* knockdown in MHCII^+^ macrophages at a 1-month time point. Knockdown is normalized to L2-siRNA^NTC^. N=3 individual mice (biological replicates) for both L2-siRNA*^Ppib^*and L2-siRNA^NTC^. Statistical significance was calculated using an unpaired two-tailed t-test (*** p<0.001). Data are presented as mean ± SD for all graphs. H. Representative image of Cy5-tagged L2-siRNA delivery to cells of the 4^th^ ventricle choroid plexus (ChP) 48 hours after injection (2 nmol in 10 μl ICV). Aqp1 marks the apical side of ChP epithelial cells. Scale bar = 100 µm (top) and 50 µm (bottom). I. scRNA-seq assessment of *Ppib* knockdown in ChP epithelial cells at a 1-month time point. Knockdown is normalized to L2-siRNA^NTC^. N=3 individual mice (biological replicates) for both L2-siRNA*^Ppib^*and L2-siRNA^NTC^. Statistical significance was calculated using an unpaired two-tailed t-test (**** p<0.0001).

Border-associated macrophages also reside in CSF outside of the brain parenchyma, primarily in the choroid plexus, meninges, and perivascular spaces, and are central players both in maintaining homeostasis and mediating CNS inflammatory responses.^19,41^ We visualized the vascular network in gray matter in three dimensions using CLARITY and light sheet microscopy and observed a high concentration of L2-siRNA within perivascular cells throughout the parenchyma (Fig. 5D). To assess the identity of these perivascular cells, we stained for macrophages based on recent work that subdivided border-associated macrophages into MHCII+ macrophages, which express genes involved in antigen presentation, and Lyve1+ scavenger macrophages, which perform a variety of essential functions including the regulation of CSF flow dynamics.^42^ We observe extensive delivery to perivascular Lyve1+ macrophages, as evidenced by localization of bright L2-siRNA puncta (Fig. 5E). MHCII+ macrophages are most abundant in the choroid plexus and meninges,^43^ and we similarly observe widespread uptake of L2-siRNA in this population (Fig. 5F). To assess knockdown in macrophages, we subclustered these populations from the L2-siRNA*^Ppib^*scRNA-seq knockdown study (Supplementary Fig. 12). While the Lyve1+ population was too sparse to evaluate, we found robust gene silencing in the MHCII+ macrophages (Fig. 5G). Lastly, we examined delivery to other cells in the choroid plexus and noted a high level of delivery to choroid plexus epithelial cells corresponding to robust gene silencing (Fig. 5H,I).

### L2-siRNA does not induce hallmarks of toxicity

To determine whether L2-siRNA is well-tolerated, we searched for potential adverse reactions, including reactive astrogliosis, neuroinflammation, and systemic organ damage. Overall, we did not observe visible signs of toxicity after surgery and every mouse injected with an L2-siRNA conjugate (Htt, Ppib, NTC) survived to their predetermined endpoint. Astrogliosis is a multi-faceted process through which astrocytes respond to damage or disease and is characterized by elevated GFAP expression, indicative of cytoskeletal hypertrophy.^44^ Immunohistochemical staining of GFAP 1 month after ICV injection did not show any differences between L2-siRNA groups (NTC, Htt) and vehicle (Fig. 6A,B). These results are consistent with Western blot quantification of GFAP in the hippocampus (Fig. 6C), collectively suggesting that L2-siRNA does not induce astrogliosis.

**Figure 6:**
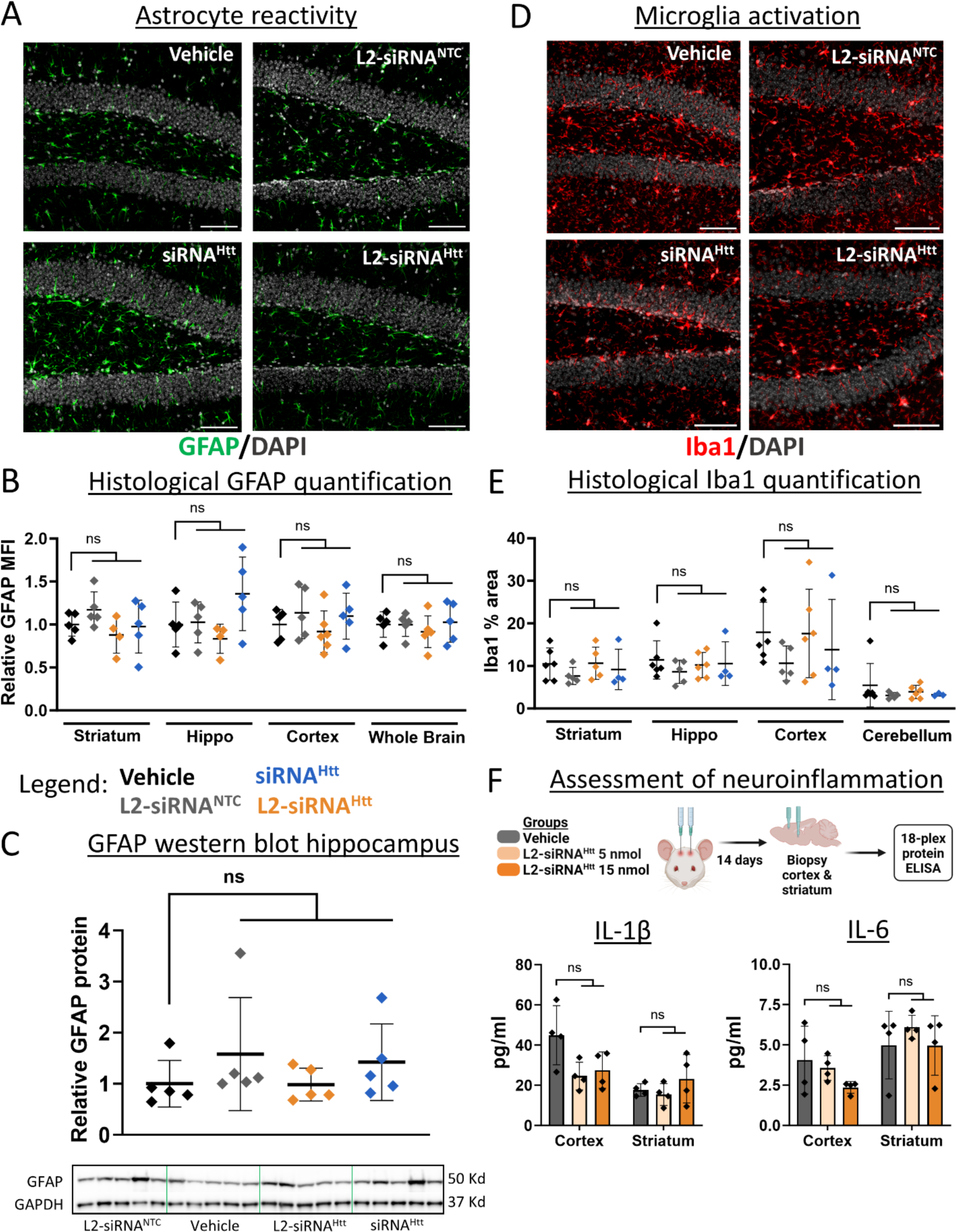
L2-siRNA does not promote CNS toxicity. A. Mice administered 15 nmol of compounds (or 0.9% saline) and stained for GFAP 1 month after ICV injection. Representative images of the hippocampal dentate gyrus are shown. Section thickness = 8 μm, scale bar = 100 μm. B. Quantification of GFAP mean fluorescence intensity (MFI) normalized to vehicle across different brain regions. Data are presented as mean ± SD, with N=4-5 mice per condition. One-way ANOVA with Bonferroni’s correction was performed for each region. C. GFAP levels measured by Western blot, normalized to GADPH housekeeping gene, and then normalized to vehicle control. Raw western blot displayed for GFAP and GAPDH. N=5 mice statistically analyzed with a one-way ANOVA and Bonferroni’s correction. D. Mice were administered 15 nmol of compounds (or 0.9% saline) and stained for Iba1 1 month after ICV injection. Representative images of the hippocampal dentate gyrus are shown. Section thickness = 8 μm, scale bar = 100 μm. E. Quantification of Iba1 protein levels normalized to vehicle. N=4-6 mice per condition. One-way ANOVA with Bonferroni’s correction was performed for each region compared to vehicle. F. Levels of inflammatory cytokines assessed two weeks after ICV injection of vehicle (0.9% saline) or L2-siRNA^Htt^ (5 nmol or 15 nmol). Additional cytokine levels are shown in Supplementary Figure 13. N=4 mice per condition. One-way ANOVA with Bonferroni’s correction was performed for each region and cytokine (ns – not significant). All data are presented as mean ± SD.

Next, we investigated whether ICV injection of L2-siRNA induced any neuroinflammatory responses. Microglia activation was assessed by Iba1 expression, indicative of a proliferative phenotype.^45^ Iba1 protein levels were unchanged after L2-siRNA injection compared to vehicle (Fig. 6D,E). To more broadly examine CNS inflammation, we utilized an ELISA panel to evaluate cortical and striatal tissue two weeks after ICV injection, at either the standard dose used in knockdown studies (15 nmol) or a lower dose (5 nmol). Notably, no difference in pro-inflammatory cytokine, chemokine, or growth factor levels were observed between vehicle and L2-siRNA^Htt^ (Fig. 6F and Supplementary Fig. 13). Consistent with prior studies showing no evidence of systemic toxicity induced by L2-siRNA after intravenous injection,^30,46^ we similarly did not observe changes in serum markers after ICV injection (Supplementary Fig. 14). Collectively, L2-siRNA did not exhibit any detectable toxicities at the doses utilized to achieve potent and durable target gene silencing, suggesting that L2-siRNA has a practical therapeutic index.

### L2-siRNA is distributed along perivascular pathways and silences *Htt* in diverse CNS regions after intrathecal delivery in rats

The CSF has several access points for drug delivery, the most common being intrathecal administration into the lumbar CSF. To more closely mimic the clinical standard, we assessed biodistribution and knockdown after intrathecal administration in rats. For this route, we were especially interested in assessing perivascular transport, as the diffusional distance from subarachnoid CSF to deep brain structures is greater in rats than in mice. We administered a bolus dose of Cy5-labeled L2-siRNA (30 nmol, ∼400 μg) through a catheter inserted via lumbar puncture into the cauda equina space.^47^ After 48 hours, we performed histology of the entire rat brain, which showed that L2-siRNA was transported along the caudal-rostral axis, where it penetrated the parenchyma from subarachnoid CSF (Fig. 7A). Robust distribution throughout all regions of the spinal cord was observed, which is expected for intrathecal delivery based on proximity to the injection site (Fig. 7B). Widespread perivascular transport throughout the brain and spinal cord was also apparent as bright red puncta. To further confirm perivascular localization, we stained vessels with rat endothelial cell antibody (RECA1), which revealed considerable transport along penetrating vessels both at the surface and in deep regions of the brain (Fig. 7C). We next assessed longer-term delivery and gene silencing of L2-siRNA^Htt^ (60 nmol, ∼800 μg) compared to a vehicle control (0.9% NaCl). The PNA assay revealed detectable levels of siRNA in every tested region, which correlated with gene silencing (Fig. 7D). Notably, L2-siRNA^Htt^ lowers *Htt* expression in all CNS regions examined, with particularly potent activity throughout the cortex, hippocampus, brainstem, and cerebellum. Collectively this evidence showcases the potential of using L2-siRNA to achieve delivery in larger species using intrathecal administration.

**Figure 7:**
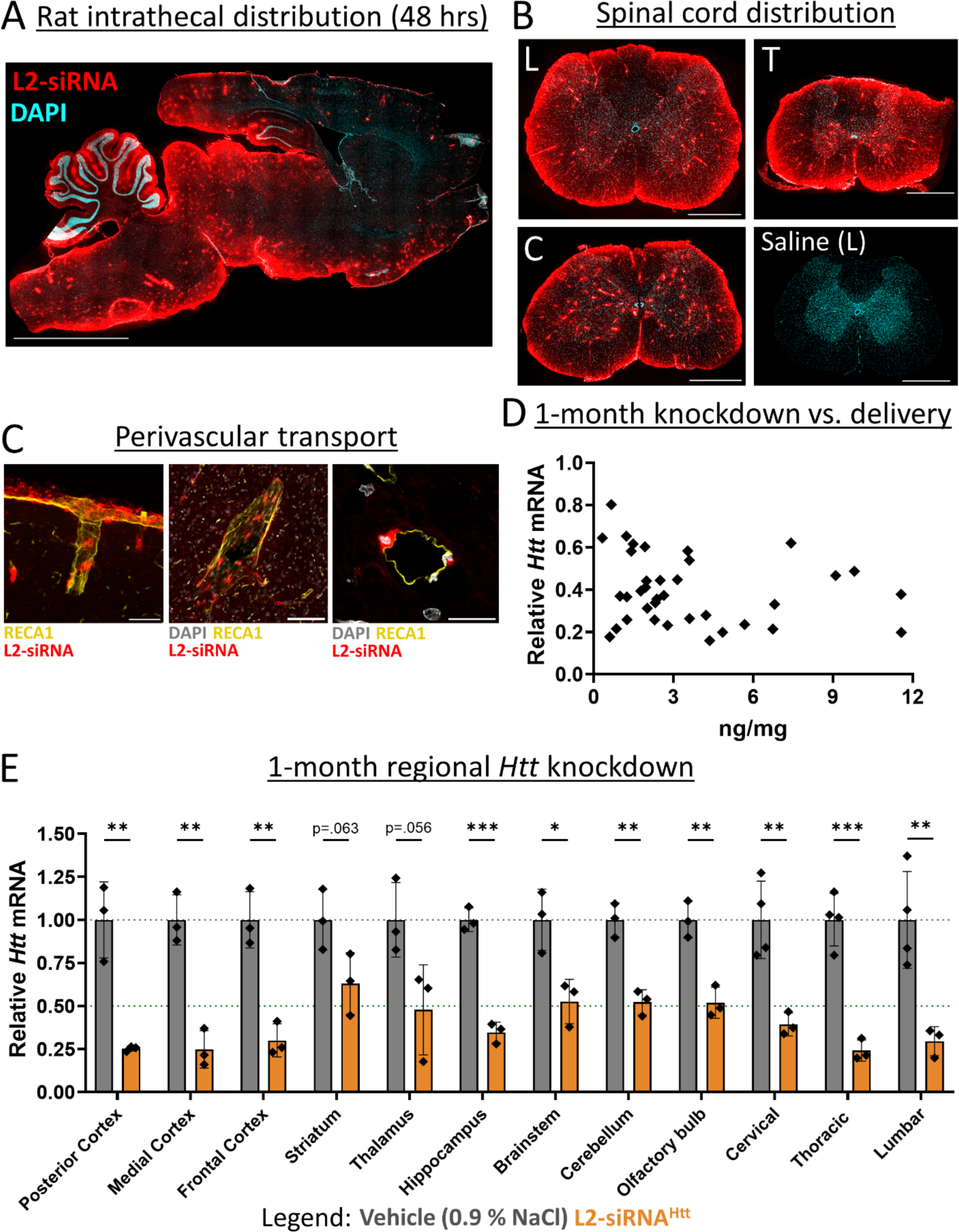
Intrathecal delivery of L2-siRNA in rats achieves broad CNS distribution and *Htt* knockdown. A. Distribution of Cy5-labeled L2-siRNA (400 μg ∼ 30 nmol) throughout the parenchyma 48-hours after intrathecal injection in rats. Scale bar = 5 mm, section thickness = 15 μm. B. Representative images of L2-siRNA delivery throughout the lumbar (L), thoracic (T), and cervical (C) regions of the spinal cord. A saline-injected rat is provided as a reference. Section thickness = 15 μm, scale bar = 1 mm. C. Perivascular transport of L2-siRNA along a representative penetrating vessel from the cortex (left image, scale bar = 50 μm) as well as around vessels in deep brain structures such as the thalamus (middle image, scale bar = 100 μm). L2-siRNA can be visualized in cells from a vessel cross section (right image, scale bar = 20 μm). D. Relationship between delivery and mRNA knockdown 1-month after intrathecal administration of L2-siRNA^Htt^ (800 μg ∼ 60 nmol). Each point represents a brain or spinal cord sample from a total of N=3 rats. E. Regional *Htt* knockdown assessed 1-month after intrathecal injection of L2-siRNA^Htt^ 800 μg ∼ 60 nmol). Each point represents an individual biological replicate (i.e., a single rat) and data are presented as mean ± SD, N=3-4. Statistical significance is calculated with unpaired two-tailed t-tests for each region (* p<0.05, ** p<0.01, *** p<0.001).

## Discussion

The clinical neurosciences are in the midst of a renaissance where, in a relatively short period of time, numerous long-intractable neurological diseases have become treatable. These advances have been mediated by a clearer understanding of the underlying disease mechanisms and the development of more-precise molecular tools. But effective treatments for some diseases, especially neurodegenerative disorders like ALS, Huntington’s disease, and Alzheimer’s disease, remain elusive. A key objective in the quest to find disease modifying therapies for these challenging disorders is delivering molecularly targeted inhibitors to specific neuroanatomical sites. Along these lines, nucleic acid delivery to the CNS remains a central challenge for the translation of gene silencing therapies, highlighting the need for rational design of nucleic acid drugs that reach deep structures of the brain after administration into the CSF. Here, we provide a detailed examination of albumin-hitchhiking L2-siRNA transport through CSF-CNS interfaces, highlighting perivascular transport and durable tissue-level silencing from a single injection targeting a diverse range of cell types in both the parenchyma and at brain borders.

CSF to brain delivery is driven by a combination of diffusion from subarachnoid/ventricular CSF and convective movement along CSF-filled perivascular spaces.^8^ Our data suggest that unconjugated siRNA disperses throughout the CNS via both pathways but is not internalized effectively by or retained within cells, as evidenced by minimal detection 48 hours after injection and rapid clearance to peripheral organs. While cholesterol conjugation has been used to promote cell uptake, our findings mirror prior studies showing that Chol-siRNA cell-association prevents effective transport across large distances.^26^ In contrast, L2-siRNA achieves a balanced cell internalization, distributing further and more homogeneously into the parenchyma from subarachnoid and perivascular CSF. These transport mechanisms were further substantiated in rats following intrathecal injection, where we observed considerable perivascular delivery throughout the spinal cord and parenchyma. We anticipate that this transport pathway will be even more critical for achieving deep, brain-wide delivery in larger mammalian species following administration into CSF.

The precise mechanisms governing drug transport through the perivascular spaces are not fully understood, but they are thought to be size dependent. For antibodies, single domain constructs exhibit greater perivascular delivery relative to full length antibodies.^8^ This suggests that relatively small nucleic acid therapies are well-suited for this transport, and indeed we observe siRNA (which are small, negatively charged linear molecules) readily accessing the entire perivascular network 2 hours after ICV delivery. Yet, the conjugation of cholesterol limits penetration into these spaces, with minimal perivascular signal beyond superficial vessels likely due to extensive interactions with cells and potentially other tissue components. A recent study showed that ASOs can be transported along the perivascular route, similar to the parent siRNA and L2-siRNA conjugate, but depth of delivery was dose dependent, with perivascular signal in deep brain structures only evident at the highest doses administered intrathecally in rats.^17^ Depending on their chemical composition (particularly the number of phosphorothioate bonds), ASOs may associate with many plasma proteins, including albumin.^48^ Along these lines, L2-siRNA association with albumin could be one of the driving forces underlying enhanced perivascular transport. We hypothesize that albumin occupancy of the L2 lipids temporarily reduces its interaction with cell membranes to enable greater dispersion throughout CSF compartments. Importantly, this interaction is reversible, and when L2-siRNA transiently dissociates from albumin, the freed lipids can facilitate efficient cell membrane binding and penetration. While further investigation of these mechanisms is warranted, our observations of L2-siRNA biodistribution has prompted us to consider perivascular delivery as a unique design principle and lends ideas through which this delivery can be enhanced in future conjugate designs.

L2-siRNA achieves potent gene inhibition comparable to other siRNA conjugates developed for CSF delivery. One such compound is the divalent siRNA, a small (∼25 kDa) and heavily phosphorothioated conjugate that exhibited potent *Htt* knockdown in mice out to 6 months following ICV delivery (40 nmol, ∼2.67X the dose used here to assess L2-siRNA knockdown).^31^ In addition, the C16 lipid conjugate developed by Alnylam® Pharmaceuticals utilizes a small hydrophobic lipid in the middle of the siRNA structure as a way to enhance cellular uptake of the siRNA without promoting micellization. While these compounds yield effective gene silencing in many regions, delivery to deep brain structures after intrathecal administration remains a considerable hurdle. Notably, C16-siRNA did not achieve knockdown in the striatum of rats, and the divalent siRNA was not effective throughout the striatum of Dorset sheep.^22,49^ Furthermore, not all cell types examined achieve effective gene silencing; for example, oligodendrocytes were not effectively silenced with C16. While gene targeting is generally assessed at the tissue level, expression of many disease-associated genes is enriched in select cell types that play outsized roles in disease progression. To address this challenge, we provide the first comprehensive examination of cell-specific gene silencing of an siRNA conjugate using scRNA-seq. While a cell-specific atlas of ASO gene silencing has previously identified cell types receptive to silencing, the analyses were only conducted in brain regions with expected gene silencing (cortex, cerebellum),^50^ whereas we investigated knockdown in cells collected from the entire brain. Further, both L2-siRNA and ASO silence a wide variety of cell types, but in a direct comparison we demonstrated that L2-siRNA^Htt^ was more potent than dose matched ASO^Htt^ across multiple brain regions, including the striatum where HD pathology manifests.

Growing evidence suggests that CSF facilitates communication between meningeal compartments and CNS parenchyma, functioning to remove waste products and transport antigens.^51^ These functions are mediated by cells residing in brain borders (of either fibroblast or myeloid origin) in both homeostatic and disease conditions.^52^ For example, recent studies implicate activation of perivascular fibroblasts as a contributing factor to neurodegeneration at early stages of Amyotrophic Lateral Sclerosis,^20^ and meningeal fibroblasts-induced fibrotic scarring has been highlighted as a potential player in the pathogenesis of multiple sclerosis.^21^ Here, we leveraged scRNA-seq to further characterize L2-siRNA targeting of CNS fibroblasts by examining gene silencing in these rare yet crucial populations residing within brain borders. Interestingly, L2-siRNA mediated gene silencing in fibroblasts spanning the arachnoid membrane, where knockdown correlated with distance from CSF; more knockdown was observed in inner arachnoid cells and arachnoid barrier cells (∼75%) than dural border cells (30%), suggesting that L2-siRNA can either cross or bypass the arachnoid barrier. We were also interested in targeting of border-associated macrophages as they are key players in a myriad of CNS disorders, including driving neurovascular dysfunction in hypertension and Alzheimer’s disease.^53,54^ One recent study identified two subtypes of macrophages, Lyve1+ and MHCII+, and demonstrated that reduced number of Lyve1+ macrophages with aging impairs CSF dynamics that can be restored with administration of macrophage colony-stimulating factor (M-CSF).^42^ MHCII+ BAMs are also implicated in neurodegeneration, for example by initiating α-synuclein-mediated neuroinflammatory responses through restimulation of CD4+ T-cells.^55^ L2-siRNA accumulates in both type of macrophages, and we observe robust gene silencing in MHCII+ macrophages. Future studies will investigate the promising therapeutic avenues of targeting these macrophages more selectively and in the context of neurodegenerative disease.

Overall, this study examined properties of lipid-siRNA conjugates that facilitate CSF to brain delivery. We performed detailed characterization of L2-siRNA transport, bulk tissue gene silencing, cell-specific activity, and toxicity. Collectively, these data suggest that L2-siRNA possesses unique properties that enhance its transport and brain-wide distribution relative to other conjugates, thus yielding a promising platform technology for silencing genes implicated in CNS disorders.

## Acknowledgments

Primary funding for this work was provided by a Chan Zuckerberg Initiative Ben Barres Early Career Acceleration Award (grant 2019-191850 to ESL), NIH grant RF1 NS130334 (to ESL and MSS), NIH grant R21 AG077807 (to ESL and CLD), and NIH R01 CA224241 (to CLD). AK and ENH were supported by the NSF Graduate Research Fellowship Program. APL was supported by the Vanderbilt Training Program in Environmental Toxicology (NIH grant T32 ES007028). Confocal microscopy was conducted in the Vanderbilt Cell Imaging Shared Resource core facility, which is supported in part by NIH grants P30 CA068485, P30 DK058404, and P30 EY008126. RNA sequencing was conducted in the Vanderbilt Technologies for Advanced Genomics core facility, which is supported in part by NIH grants 5UL1 RR024975, P30 CA068485, P30 EY008126, UL1 TR002243, and G20 RR030956. Whole slide imaging for toxicity analysis was performed in the Digital Histology Shared Resource at Vanderbilt University Medical Center. FACS sorting experiments were performed in the VMC Flow Cytometry Shared Resource, which is supported by the Vanderbilt Ingram Cancer Center (P30 CA68485) and the Vanderbilt Digestive Disease Research Center (DK058404). All LC-MS analyses were performed in the Mass Spectrometry Research Center at Vanderbilt University. We would also like to thank Karen Thompson and the Translational Pathology Shared Resource for assistance with cryosectioning (NCI/NIH Cancer Center Support Grant P30CA068485). Figures 1A,B; 2A; 4A,D,F; 5A,B; 7A; and Supplementary Figs. 10A,B; 11A; 12A,C; and 14A were created with BioRender.com.

## Author contributions

AGS, CLD, and ESL conceived the project. AGS performed the majority of experiments with assistance from KRS, AMA, AS, KAK, RPC, JCP, APL, WTF, LVA, ENH, AP, MC, and DLM. NF performed all syntheses. AK performed all computational analyses. KMS and TMM supervised and assisted with rat intrathecal injections. KCV oversaw analyses of conjugate interactions in CSF. MSS provided de-identified CSF samples and oversaw lightsheet imaging and mouse surgical procedures. CLD and ESL oversaw all work. AGS, CLD, and ESL wrote the original manuscript draft. All authors reviewed and edited the manuscript.

## Data availability

Raw sequencing data and processed Seurat objects are available at Array Express under accession number E-MTAB-13964. Code for reproducing the single-cell RNA sequencing analysis is available upon request.

## Ethics declaration

The authors declare no competing interests.

## Methods

### Synthesis of oligonucleotides

Oligonucleotide syntheses were performed using standard solid-phase chemistry with a MerMade 12 automated RNA synthesizer (BioAutomation) on controlled pore glass with a universal support (1 or 10 μmol scale, 1000Å pore), using 2’-F and 2’-OMe phosphoramidites with standard protecting groups (Glen Research). 5′-(E)-Vinylphosphonate (VP) was incorporated on the 5’ terminus of the antisense strand using POM-vinyl phosphonate 2’-OMe-uridine CE-phosphoroamidite (LGC genomics). All strands were grown on the universal support except for Cy5-labelled oligos, which were synthesized on Cy5-functionalized 1000Å CPG (Glen Research). Amidites were all dissolved at 0.1 M in anhydrous acetonitrile, except for 2′-OMe uridine which requires 20% dimethylformamide as a co-solvent. All nucleic acid sequences are listed in Supplementary Table 1.

### Cleavage and deprotection of oligonucleotides

Unconjugated and conjugated sense strands were cleaved and deprotected with a 1:1 solution of 28%– 30% ammonium hydroxide and 40% aqueous methylamine (AMA) for 2 hours at room temperature. Cy5-labeled oligonucleotides were cleaved in 28–30% ammonium hydroxide at room temperature for 20 hours. The VP-containing antisense strands were cleaved and deprotected by treating the CPG with a 3% diethylamine (DEA) solution in 28–30% ammonium hydroxide (20 hours, 35°C).

### Purification and characterization of oligonucleotides

After cleavage and deprotection, oligonucleotides were dried under vacuum to remove solvents (Savant SpeedVac SPD 120 Vacuum Concentrator, ThermoFisher). Pellets were resuspended in water and purified on a Waters 1525 EF HPLC system equipped with a Clarity Oligo-RP column (Phenomenex) under a linear gradient [60% mobile phase A (50 mM triethylammonium acetate in water) to 90% mobile phase B (methanol)]. Cy5-labeled and unconjugated (DMT-on) sense strands were first desalted using Gel-Pak column (Glen Research) followed by chromatography under a linear gradient (85% to 40% mobile phase A). Oligonucleotide fractions were dried, resuspended in nuclease free water, sterile filtered, and lyophilized. DMT protecting group was removed from the purified, dried, unconjugated strand using 20% acetic acid for 1 hour at room temperature, followed by desalting.

Antisense strands were purified over a 10 x 150 mm Source 15Q anion-exchange column (Cytiva) using a gradient of sodium perchlorate. Buffer A consisted of 10 mM sodium acetate in 20% acetonitrile and Buffer B consisted of 1 M sodium perchlorate in 20% acetonitrile. The run conditions were 90% to 70% buffer A over 30 minutes at a flow rate of 5 ml/min. Purified oligonucleotides were desalted, sterile filtered, and lyophilized.

The identity of oligonucleotides was verified by Liquid Chromatography-Mass Spectrometry (LC-MS, ThermoFisher LTQ Orbitrap XL Linear Ion Trap Mass Spectrometer). LC-MS was performed using a Waters XBridge Oligonucleotide BEH C18 Column under a linear gradient [85% phase A (16.3 mM triethylamine – 400 mM hexafluoroisopropanol) to 90% phase B (methanol)] run for 10 minutes at 45°C.

### *In vitro* assessment of carrier-free oligonucleotide uptake

Uptake propensity of siRNA conjugates was assessed by treating N2a neuroblastoma cells with Cy5 labeled compounds and measuring fluorescence by flow cytometry. In brief, N2a cells (<P21) were seeded at 100,000 cells/ml onto uncoated 24-well plates and allowed to adhere overnight. The cells were then treated with siRNA-Chol, L2-siRNA, or free siRNA in serum-free Opti-MEM at 60 nM (Cy5 concentration) for 2 hours. Unbound compound was removed with two DPBS-/-washes and then the cells were dissociated using Accutase (Sigma). The collected cells were pelleted and resuspended in flow cytometry buffer (0.5% BSA in DPBS without calcium and magnesium) to run on a Guava EasyCyte (Luminex) flow cytometer. The analysis was performed in FlowJo™ v10.8 Software (BD Life Sciences) to gate single cells (from debris and doublets) and measure geometric mean fluorescence intensity from over 2,000 cellular events.

### *In vitro* assessment of carrier-free gene silencing

*In vitro* gene silencing of oligonucleotide conjugates was assessed by carrier-free reverse transfection. Wells were prepared with 250 nM of siRNA in Opti-MEM, then 75,000 N2a cells were added to each well (24-well plate). After 24 hours, an equal volume (500 μl) of full serum DMEM (10% FBS) was added to each well and after an additional 24 hours, cells were harvested for RT-qPCR.

### Animal husbandry

Adult C57BL/6J male mice were ordered from Jackson Laboratory and used between 10-16 weeks of age. Mice were housed continuously in an environmentally controlled facility in a 12-hour light/dark cycle with *ad libitum* access to food and water. All mouse protocols were approved by the Institutional Animal Care and Use Committee (IACUC) at Vanderbilt University. One-month rat intrathecal studies were performed in male Sprague Dawley rats at Vanderbilt and approved by Vanderbilt’s IACUC. All 48-hour rat studies were conducted in 69-84 day old Sprague Dawley rats (male and female) and approved by the IACUC at Washington University in St. Louis.

### Intracerebroventricular (ICV) injections and euthanasia

siRNA duplexes were annealed in 0.9% sterile saline by heating to 95°C and gradually cooling to 4°C on a thermocycler. On the day of injection, the compounds were concentrated to either 1.5 mM (15 nmol; for gene silencing) or 1 mM (10 nmol; Cy5-tagged siRNA; for biodistribution) using a 3K Amicon Ultra spin filter (UFC500324). The concentrations were measured by absorbance (260 nm) on a NanoQuant Tecan plate reader and adjusted as necessary by adding saline.

Mice were anesthetized with isoflurane and mounted on the stereotactic rig where they receive continuous isoflurane for the duration of the surgery. Eye ointment (Bausch + Lomb) was administered to prevent drying and then the scalp was sanitized with betadine and 70% ethanol, alternating three times each. The scalp was opened with a midline incision and hydrogen peroxide was applied to expose bregma. Injection coordinates, as distance from bregma, were ± 1 medial-lateral, -2.3 dorsal-ventral, and -.2 anterior-posterior. Two holes were drilled through the skull at these coordinates for bilateral injection. The syringe (Hamilton Model 701, blunt 30 G) was brought to these coordinates and slowly lowered into the ventricle. Injections were performed at 1 µl/min for a total of 5 µl per ventricle. To minimize backflow, the needle was left in the ventricle for an additional 5 minutes prior to gradual retraction. The scalp was then sutured shut, and mice were monitored for their recovery. To maintain body temperature, mice were placed on a heating pad (37°C) during and after surgery. For analgesia, mice received an intraperitoneal injection of Ketoprofen (5-10 mg/kg Ketofen, Zoetis) prior to surgery and daily for 72 hours post-operation.

At the terminal timepoint, mice were euthanized by ketamine (450 mg/kg)/xylazine (50 mg/kg) overdose and transcardially perfused with cold heparinized (10 U/ml) DPBS-/-to remove blood cells from the vasculature. For flow cytometry studies, the right hemisphere was extracted into DPBS and processed into single cells as described below. For gene silencing studies, brains were extracted after perfusion and cut into 1 mm slices using a sagittal brain matrix. Biopsy punches (2 mm) were taken from different brain regions (hippocampus, striatum, cerebellum, posterior cortex, brainstem) and stored in RNAlater (Thermo AM7020) for downstream analyses. Spinal cords were extracted by extrusion with HBSS and segmented into cervical, thoracic, and lumbar regions using the characteristic enlargements as a guide. For biodistribution studies, mice were further perfused with 4% paraformaldehyde (PFA), and then brains and spinal cords were extracted, immersion fixed in 4% PFA overnight at 4°C, and subjected to further downstream histological processing. Organs were also harvested and Cy5 fluorescence was measured by IVIS Lumina III imaging (Caliper Life Science, Hopkinton, MA).

### Rat intrathecal injections and euthanasia

Rat intrathecal administration of siRNA conjugates was performed as described^47^ by inserting a catheter into the cauda equina space and injecting a bolus of L2-siRNA. To prepare for surgery, rats were anesthetized with isoflurane, shaved, and disinfected by alternating betadine and 70% ethanol for three cycles. After locating the injection site between L6 and L5, an incision was made in the skin and connective tissue removed by blunt dissection. Local anesthetic (50 μl of 0.5% bupivacaine) was applied intramuscular near the spinal column and allowed to settle before inserting the guide cannula (19G blunted needle) into the thecal sac. Next, a catheter was threaded through the guide cannula and into the cauda equina space. Successful placement of the catheter was confirmed by visualization of CSF backflow. A bolus dose of L2-siRNA (30 μl, ∼1 μl/second) was administered, followed by 40 μl of saline. The catheter was then heat-sealed and the incision closed with staples or sutures. Rats then received Carprofen (5 mg/kg intraperitoneal) and were monitored for recovery on a warming pad. For 48-hour biodistribution studies, rats were euthanized with Fatal Plus (200 mg/kg) and perfused with heparinized PBS (10 U/ml). For 1-month studies, rats were perfused with DPBS (-/-) under isoflurane.

### Peptide nucleic acid (PNA) hybridization assay

We adapted a previously described PNA assay to quantify absolute siRNA delivery after ICV administration in mice and intrathecal in rats.^56,57^ Conceptually, the siRNA duplex is denatured into single strands, and a Cy3-PNA probe with complete complementarity to the antisense strand forms a new PNA-antisense duplex. When run through an anion-exchange column, the PNA-RNA duplexes elute in a distinct peak, and the area under the curve can be related to mass of siRNA using a standard curve where known amounts of siRNA are doped into tissue homogenates from untreated mice or rats. Data are reported as mass of antisense strand per mass of tissue.

To prepare homogenates, tissue biopsy punches were removed from RNAlater, placed in 300 μl homogenization buffer (Thermo QS0518) plus proteinase-K (Thermo QS0511, 1:100), and disrupted using a Tissuelyzer 2.0 for 5 minutes at 30 Hz. Following a 1-hour incubation at 65°C, the samples were spun down at 15,000 G for 15 minutes and the supernatant was collected for storage at -80°C. The standard curve was prepared at the same time as the homogenates, with a maximum of 10,000 fmol and minimum of 156.25 fmol by 1:2 serial dilution. A new standard curve was prepared for each conjugate type and siRNA sequence.

Samples were thawed and sodium dodecyl sulfate (component of homogenization buffer) was precipitated from 200 μl of homogenate with 20 μl of 3M potassium chloride and centrifuged at 4,000 × g for 15 min. The supernatant was collected and centrifuged at the same speed for an additional 5 minutes to ensure complete removal of the precipitate. If samples required a dilution to fall within the standard curve, they were diluted to 200 μl in homogenization buffer prior to precipitation. Next, 150 μl of supernatant was transferred to a screw cap tube, where 100 μl of hybridization buffer (50 mM Tris, 10% ACN, pH 8.8) and 2 μl of 5 μM PNA probe (∼10 pmol/ 150 ul of sample, PNA bio) were added. The probe was annealed to the antisense strand by heating to 90°C and then 50°C for 15 minutes each. The samples were then run through a DNAPac PA100 anion-exchange column (Thermo Fisher Scientific) on an iSeries LC equipped with RF-20A fluorescence detector (Shimadzu). Mobile phases consisted of buffer A (50% acetonitrile and 50% 25 mM Tris–HCl, pH 8.5; 1 mM ethylenediaminetetraacetate in water) and buffer B (800 mM sodium perchlorate in buffer A), and a gradient was obtained as follows: 10% buffer B within 4 minutes, 50% buffer B for 1 minutes and 50% to 100% buffer B within 5 minutes.^58^ The final mass of siRNA was calculated using the area under the curve of Cy3 fluorescence from a standard curve of known quantities of siRNA or L2-siRNA spiked into untreated tissue homogenates.

### RT-qPCR

RT-qPCR methodology was employed to determine mRNA silencing *in vitro* and *in vivo*. For *in vitro* studies, cells were lysed in RLT buffer plus β-mercaptoethanol (1:100) and RNA was extracted using an RNeasy plus mini kit (Qiagen 74134). Reverse transcription into cDNA was performed according to iScript manufacturer instructions (iScript cDNA synthesis kit, Biorad 1708891). Gene expression was measured by Taqman qPCR on the cDNA using 20 μl reactions, run on a Biorad CFX96, and analyzed in CFX Maestro software.

For *in vivo* studies, brain homogenates were prepared in 350 µl of RLT buffer plus β-mercaptoethanol and processed with 5 mm stainless steel beads (Qiagen cat. No. 69989) for 5 minutes at 30 Hz (TissueLyser II). RNA was then extracted using an RNeasy plus micro kit (Qiagen 74034) according to manufacturer instructions. RNA was eluted in 14 μl RNAse-free water and the concentration and purity were measured by 230/260/280 nm absorbance on a Nanodrop 2000c spectrophotometer. Last, qPCR was performed by preparing a 10 µl reaction mixture composed of 2X master mix, water, Taqman probes, and cDNA sample. Taqman probes were Mm00478295_m1 (*Ppib*), Mm01213820_m1 (*Htt*), Rn00577462_m1 (Htt), Rn03302274_m1 (*Ppib*), Mm02619580_g1 (*Actb*). *In vivo* samples were run on a Quant 12k flex in triplicate (2 min @ 50^◦^C, 10 min @ 95^◦^C, then cycle 15 seconds @ 95^◦^C and 1 min @ 60^◦^C). All samples were analyzed according to standard ΔΔCt methodology. Each sample is normalized to *Ppib* as a housekeeping gene unless *Ppib* is the target of the siRNA, in which case *Actb* is the housekeeping gene. Conventional RT-qPCR controls (no-template control and no reverse transcriptase control) were run on every plate and did not amplify.

### Western blotting

Brain homogenates were prepared in freshly made lysis buffer containing 10 mM HEPES (Sigma, adjusted to pH 7.2), 250 mM sucrose (Sigma), 1 mM EDTA, 1 mM sodium fluoride, 1 mM sodium orthovanadate, and 1 protease inhibitor tablet (Roche 11836170001). 75 µl of buffer was added to each biopsy punch, and samples were homogenized three times for 10 seconds with a handheld homogenizer. Samples were cooled on ice for 30 seconds between homogenizations to prevent protein degradation. All homogenates were then sonicated (handheld) for 20 seconds to liberate nuclear proteins, and samples stored at -80°C until further analysis. Bicinchoninic acid (BCA) assay (Thermo 23225) was used to quantify protein and normalize loading of each sample. In brief, 10 µl of sample was added to each well in duplicate followed by the addition of 200 µl of substrate. The plate was incubated for 30 minutes at 37°C in the dark. Absorbance was measured at 562 nm and protein was quantified from a bovine serum albumin standard curve.

To prepare samples for Western blotting, 10 µg of each sample was diluted with TBS in XT sample buffer (Biorad 1610791) plus 5 mM dithiothreitol (dtt) and beta-mercaptoethanol (1:10) and heated for 5 minutes at 95°C to fully denature proteins. Samples were loaded on a 3-8% tris-acetate gel (Biorad 3450131) in Tricine running buffer (Biorad 1610790) or Novex tricine SDS running buffer (LC1675). The gel was run at 120-150V until the loading dye ran off the bottom. Next, the gel was transferred to a nitrocellulose Midi membrane (Biorad 1704159) using a Trans-Blot Turbo (Biorad). This membrane was then blocked for 1 hour in 5% non-fat milk (Bob’s red mill) diluted in TBS. Primary antibodies were added overnight in 5% milk in TBS + 0.1% tween-20 (TBST): huntingtin (Clone D7F7, Cell Signaling 5656, 1:1,000), beta-actin (Clone 13E5, Cell signaling 4970, 1:2,000), GFAP (Clone D1F4Q, Cell signaling 12389, 1:2,000). After rocking at 4°C overnight, the membrane was washed four times for 10 minutes with TBST. The HRP-conjugate secondary antibody (Abcam ab6721) was diluted 1:20,000 in TBST and applied at room temperature for 1 hour. When applicable, GAPDH (HRP-60004 Proteintech, 1:3,000) was added with the secondary. After washing thrice with TBST and once with TBS for 10 minutes, the HRP substrate was added to visualize the bands. For huntingtin detection, super signal west femto substrate (Thermo 34095) was used, while Clarity Western ESC (Biorad 1705060) was used for beta actin detection. Blots were imaged for chemiluminescence on a Bio-rad ChemiDoc MP imaging system and analyzed in ImageLab to measure total band intensity.

### Histology

To prepare fixed brains for frozen sections, sucrose gradients were performed for cryoprotection by immersion in 15% sucrose for 24 hours (or until the sample sinks), followed by immersion in 30% sucrose for an additional day. The hemispheres were then embedded in Epredia Neg-50 medium, sectioned to 30 μm on a cryostat, and stored at -80°C until staining.

Antibody staining was performed as follows. First, samples were thawed to room temperature and a barrier was drawn using a hydrophobic pen to localize the staining reagents on the sample. Samples were washed in DPBS-/-with 0.3% Triton-X 100 (PBST), followed by blocking for 1 hour at room temperature in PBST plus 5% donkey serum (Sigma D9663). Slides were then incubated either 2 hours at room temperature or overnight at 4°C with the primary antibody diluted in DPBS containing 1% BSA and 0.5% Triton-X 100. Primary antibodies include CD31 (1:100, BD biosciences 550539), Lyve1 (1:200, R&D AF2125), AQP4-488 (1:500 for overnight or 1:200 for 2 hours, Abcam ab284135), Glut1-PE (1:300, Abcam ab209449), AQP1 (1:200, Abcam ab168387). The sections were then washed three times in PBST for 5 minutes, followed by a 1-hour room temperature incubation in the appropriate secondary antibody (1:500 dilution in DPBS containing 1% BSA and 0.5% Triton-X 100). Sections were washed, then incubated with DAPI (1:5,000-1:10,000, 5 mg/ml stock, Thermo D1306) for 10 min, washed again in DPBS and then mounted under a coverslip with ProLong gold antifade reagent. Imaging was performed either on a Leica epifluorescence microscope (tiled images of entire brain) or a Zeiss LSM 710 confocal microscope (higher magnification of select regions). Image processing was performed using Fiji software. For samples where antibody staining was not required (i.e. assessing Cy5-tagged siRNA signal), slides were washed twice with DPBS and then mounted with ProLong gold plus DAPI.

Toxicity immunohistochemistry was performed on paraffin embedded sections using the Epredia Autostainer 360. Sections were baked at 60^◦^C for 1 hour then deparaffinized and rehydrated through xylene and ethanol steps. Antigen retrieval was performed using the Epredia PT module in citrate buffer (pH 6.0) at 97^◦^C. Slides were then transferred to the autostainer and blocked with Dako Protein Block (X0909) and Flourescent block (ThermoFisher 37565). Primary antibodies were diluted in Dako protein block and incubated on the slides for 1 hour at room temperature. Astrocytes were labeled with anti-GFAP (1:1000, Dako Z0334) and microglia with anti-Iba1 (1:500, Wako 019-19741). Slides were then washed and incubated in Alexaflour 647-conjugated secondary antibody (1:1000, ThermoFisher A21245) for 1 hour at room temperature. Sections were mounted with prolong gold with DAPI (Invitrogen P36931) and imaged using the Aperio Versa 200 slide scanner. All quantification was done using ImageJ.

### Tissue clearing

Mouse brains or spinal cords were incubated for 3-5 days in 4% PFA and the right hemispheres were embedded in CLARITY polyacrylamide hydrogel (4% Acrylamide, 0.05% Bis-Acrylamide, 0.25% temperature-triggering initiator VA-044 in 0.1 M PBS). To allow time for the polyacrylamide to permeate, the tissue remained in unpolymerized solution at 4°C for 2 weeks. Tubes were then moved to a 37°C water bath for 4 hours to activate the VA-044 acrylamide crosslinker and polymerize the hydrogel. After the polymerization, tissue was cleared passively with a clearing solution (200 mM boric acid, 4% w/v SDS, pH 8.5) at 37°C shaking in a humidified incubator for 4-8 weeks. To stain for vasculature, the sample was washed overnight in PBS (0.1 M) with 0.1% Triton-X and then incubated in the same buffer containing lectin-fluorescein (0.5 mg/ml, FL-1171, Vector Laboratories), overnight for spinal cords and two days for brains. The same wash was repeated overnight, and afterwards the samples were incubated in 68% Thiodiethanol (TDE) overnight for refractive index matching (1.33). The samples were imaged on a light-sheet Z1 microscope (Zeiss) using 20x objective illumination and processed with Imaris software (version 9.9.0, Bitplane, USA).

### Fast protein liquid chromatography (FPLC)

FPLC was used to assess binding of different compounds to albumin in human CSF. Cy5-labeled conjugates or free siRNA (1 µM) were mixed with 300 µl post-mortem human CSF or human serum albumin (7.5 μM). After 30 minutes of incubation at 37°C, the volume was then brought to 1 ml in running buffer (10 mM Tris-HCl, 0.15 M NaCl, 0.2% NaN3), filtered (0.22 μm, Millipore UFC30GV00), and fractionated by size-exclusion through three tandem superdex 200 increase columns on an Akta Pure FPLC (GE Healthcare). The siRNA content of each fraction was measured with a plate reader (Biotek Synergy H1) for Cy5 fluorescence.

### Flow cytometry

Harvested brain tissue was converted to single cell suspensions using the papain-based mouse adult brain dissociation kit from Miltenyi Biotec (130-107-677), with appropriate steps for myelin removal and red blood cell lysis according to the manufacturer’s instructions. Cells were then FcR-blocked for 10 minutes on ice (10 µl per sample, Miltenyi 130-092-575). The cells from each brain were then split into 4 experimental samples – one for each cell-specific antibody cocktail: Thy1 for neurons (1:5,000; Biotechne FAB7335P), ACSA2 for astrocytes (1:2,000; Miltenyi 130-123-284), CD11b for immune cells (1:2,000; BD biosciences 561689), and CD31 for endothelial cells (1:2,000; eBioscience 17-0311-80). Each population was additionally stained with O1 (1:100, R&D systems FAB1327G), a marker for oligodendrocytes, which are excluded from analysis. After washing, cells were resuspended in DPBS-/- with 0.5% BSA and DAPI (1:10,000, 5 mg/ml stock) and run on an Amnis CellStream flow cytometer. 25,000 cellular events were recorded. The gating scheme to identify Cy5+ cells leveraged fluorescence-minus-one (FMO) controls, which are samples stained with identical antibodies to the experimental groups but lack Cy5 signal (originating from an uninjected mouse). Single color compensation controls were run for each fluorophore. To validate ACSA2 gating of astrocytes, cells were sorted on a 4-laser BD FACSAria III and RT-qPCR was performed as described using taqman probes to show enrichment of astrocytes (Mm01253033_m1) and depletion of endothelial cells (Mm00727012_s1).

### Multiplexed analysis of cytokines

Mice were administered 15 or 5 nmol of L2-siRNA by ICV injection as previously described. Two weeks after ICV injection, the cortex and striatum were biopsy punched, flash frozen, and stored at -80 °C. To prepare homogenates, samples were thawed in cell lysis buffer (Biorad 171304006M) and homogenized three times for 10 seconds with a handheld homogenizer, with 30 second intervals on ice between homogenizations. The solution was then sonicated for 30 seconds and centrifuged at max speed for 10 minutes at 4°C. Supernatant was collected and total protein was quantified by BCA assay as previously described. The samples were then diluted to 1 mg/ml and processed by Eve Technologies using the Mouse High Sensitive 18-Plex Discovery Assay according to the manufacturer’s protocol (Millipore Sigma).

### Single-cell RNA sequencing (scRNA-seq) sample preparation

Mice were administered 15 nmol of L2-siRNA by ICV injection as previously described. The siRNA sequence was targeted to *Ppib* or a non-targeting control. After 1 month, mice were perfused with heparinized (10 U/ml) DPBS and a single cell suspension of the brain was prepared using the adult brain dissociation kit as previously described. CD11b positive and negative cells were isolated via MACS sorting with Cd11b-coated microbeads (Miltenyi 130-097-142) according to the manufacturer’s instructions. After 2 washes with cell-suspension buffer, cells were counted on a hemocytometer and diluted to target 2,000 (Cd11b^pos^) or 20,000 (Cd11b^neg^) cells for encapsulation. Unencapsulated cells were used for RT-qPCR as previously described. The single cell libraries were prepared using the PIPseq T2 (for Cd11b^pos^) or v4.0 PIPseq T20 kits (for Cd11b^neg^) following manufacturer specifications. Reads were generated at a depth of 40 million reads for CD11b^pos^ and 400 million read for CD11b^neg^ from an Illumina NovaSeq6000 PE150 sequencing run, yielding a final read depth of 20K reads per cell.

### scRNA-seq data processing

Reads were processed using the PIPseeker alignment algorithm (v02.01.04). Counts matrices were then processed using standard techniques in Seurat (v4). Datasets were merged without batch correction, and cells were filtered based on number of features, keeping cells with 800–4,000 uniquely expressed genes for samples processed with T2 kits and 800-10,000 for samples processed with T20 kits. Counts were then normalized, and the dataset was reduced to the top 2,000 variable genes. The data were then scaled, and dimension reduction (PCA and UMAP) was performed based on the first 50 principal components to visualize the data. Cells were clustered using the Louvain algorithm and annotated based on standard marker genes according to the mouse brain atlas.^59^ Average *Ppib* expression levels were calculated using the AverageExpression() function in Seurat and were reported per cluster per biological replicate. For barplots, expression levels were normalized to L2-siRNA^NTC^ control.

