## Supplemental information for "Lipid-siRNA conjugate accesses perivascular transport and achieves durable knockdown throughout the central nervous system"

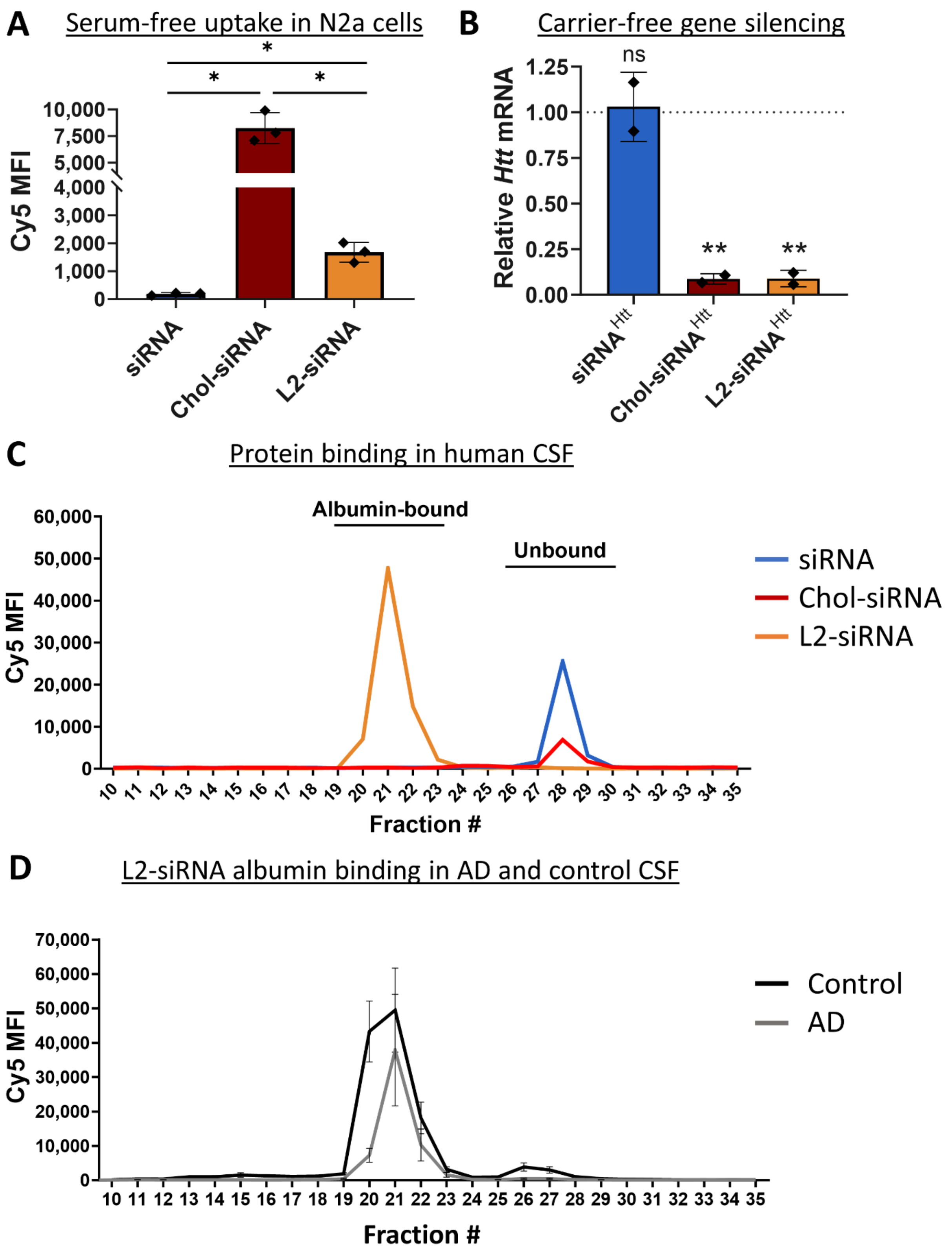

A
Serum-free uptake in N2a cells
Cy5 MFI
siRNA
Chol-siRNA
L2-siRNA
B
Carrier-free gene silencing
Relative Htt mRNA
ns
**
siRNA^Htt
Chol-siRNA^Htt
L2-siRNA^Htt
C
Protein binding in human CSF
Albumin-bound
Unbound
Cy5 MFI
Fraction #
siRNA
Chol-siRNA
L2-siRNA
D
L2-siRNA albumin binding in AD and control CSF
Cy5 MFI
Fraction #
Control
AD

**Supplementary Figure 1: Physicochemical properties of siRNA conjugates**

A. Cell uptake was evaluated by flow cytometry in N2a cells after a 2-hour incubation with unconjugated siRNA, Chol-siRNA, or L2-siRNA (60 nM). Mean fluorescence intensity (MFI) is reported from three independent experiments, where the bars represent mean ± SD. One-way ANOVA without assuming equal SD was performed with Dunnett's T3 correction for multiple comparisons (*$p<0.05$).

B. Carrier-free *Htt* knockdown in N2a cells assessed by RT-qPCR after a 48-hour incubation with $siRNA^{Htt}$, $Chol\text{-}siRNA^{Htt}$, or $L2\text{-}siRNA^{Htt}$ normalized to $L2\text{-}siRNA^{NTC}$ (represented by dotted line at Y=1.0). Data reported from two independent experiments, each normalized to $L2\text{-}siRNA^{NTC}$, with bars representing mean ± SD. One-way ANOVA compared to $L2\text{-}siRNA^{NTC}$ with Bonferroni's multiple comparison correction (ns – not significant, **$p<0.01$).

C. Albumin-binding properties of siRNA conjugates in human CSF. Cy5-labeled siRNA, Chol-siRNA, or L2-siRNA (1 μM) mixed with 300 μl of human CSF was analyzed using fast protein liquid chromatography (FPLC). Only L2-siRNA elutes in known albumin-containing fractions. Same N=1 CSF sample used for each condition.

D. L2-siRNA associates with albumin in CSF from patients with Alzheimer's Disease (AD) as well as those without neurodegeneration (N=3 control and AD samples). Error bars represent SD.

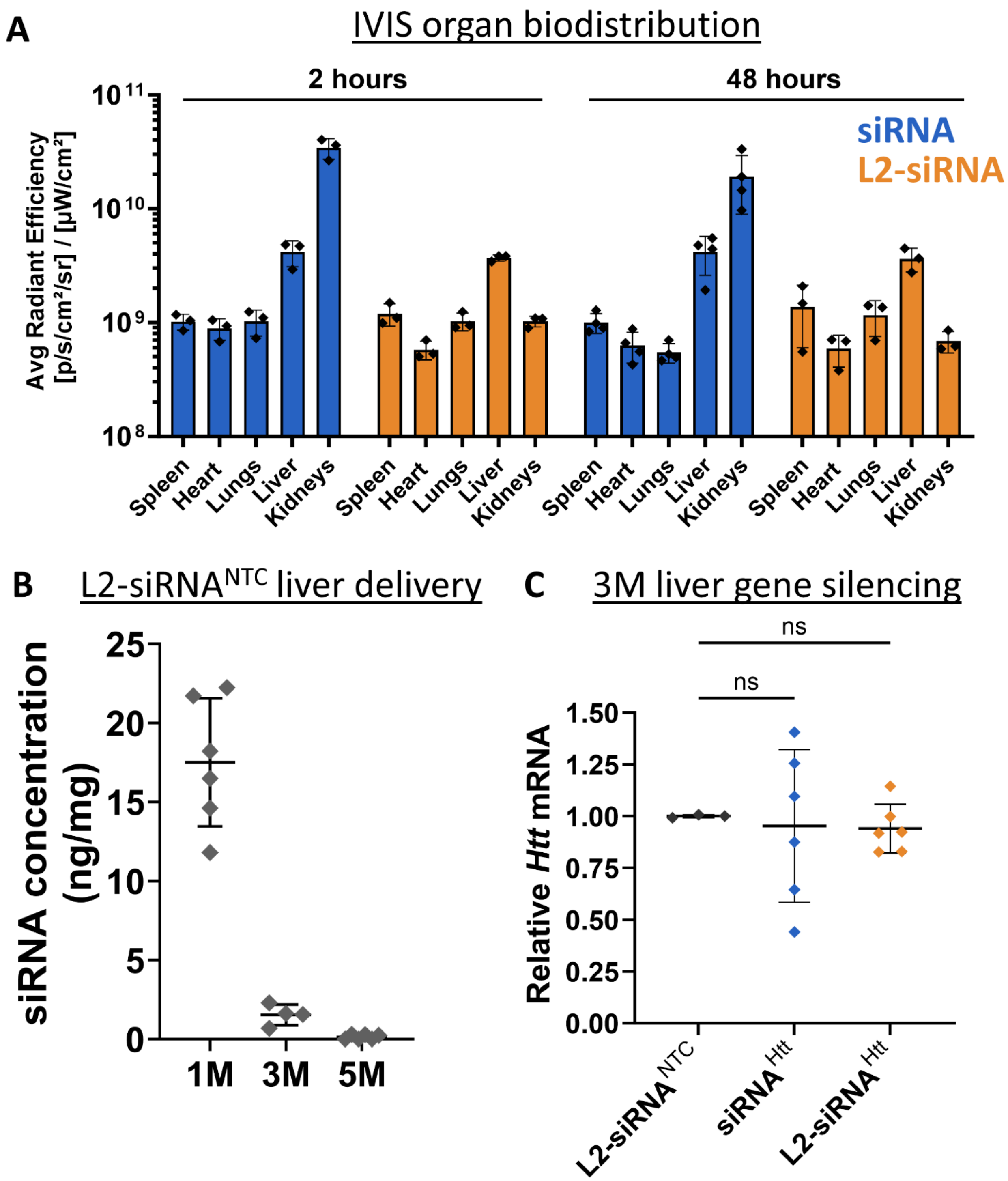
A
IVIS organ biodistribution
2 hours
48 hours
siRNA
L2-siRNA
Avg Radiant Efficiency [p/s/cm²/sr] / [µW/cm²]
10^11
10^10
10^9
10^8
Spleen
Heart
Lungs
Liver
Kidneys
B
L2-siRNA^NTC liver delivery
siRNA concentration (ng/mg)
25
20
15
10
5
0
1M
3M
5M
C
3M liver gene silencing
ns
ns
Relative Htt mRNA
1.50
1.25
1.00
0.75
0.50
0.25
0.00
L2-siRNA^NTC
siRNA^Htt
L2-siRNA^Htt

**Supplementary Figure 2: Clearance to peripheral organs**

A. Accumulation in peripheral organs assessed by In Vivo Imaging System (IVIS) 2 and 48 hours after ICV injection of Cy5-labeled siRNA or L2-siRNA (7.5-10 nmol). Average radiant efficiency of Cy5 fluorescence is reported. N=3-4 mice.

B. L2-siRNA$^{NTC}$ liver delivery 1, 3 and 5 months (M) after ICV injection (15 nmol) measured with the PNA assay. Values below the limit of detection were plotted as 0. N=4-6 mice.

C. *Htt* mRNA expression levels in the liver as measured by RT-qPCR 3 months after ICV injection (15 nmol). Each point represents an individual mouse (N=3-6). Significance was calculated as a one-way ANOVA compared to L2-siRNA$^{NTC}$ with Bonferroni's correction for multiple comparisons. Data presented as mean ± SD in every graph (ns - not significant).

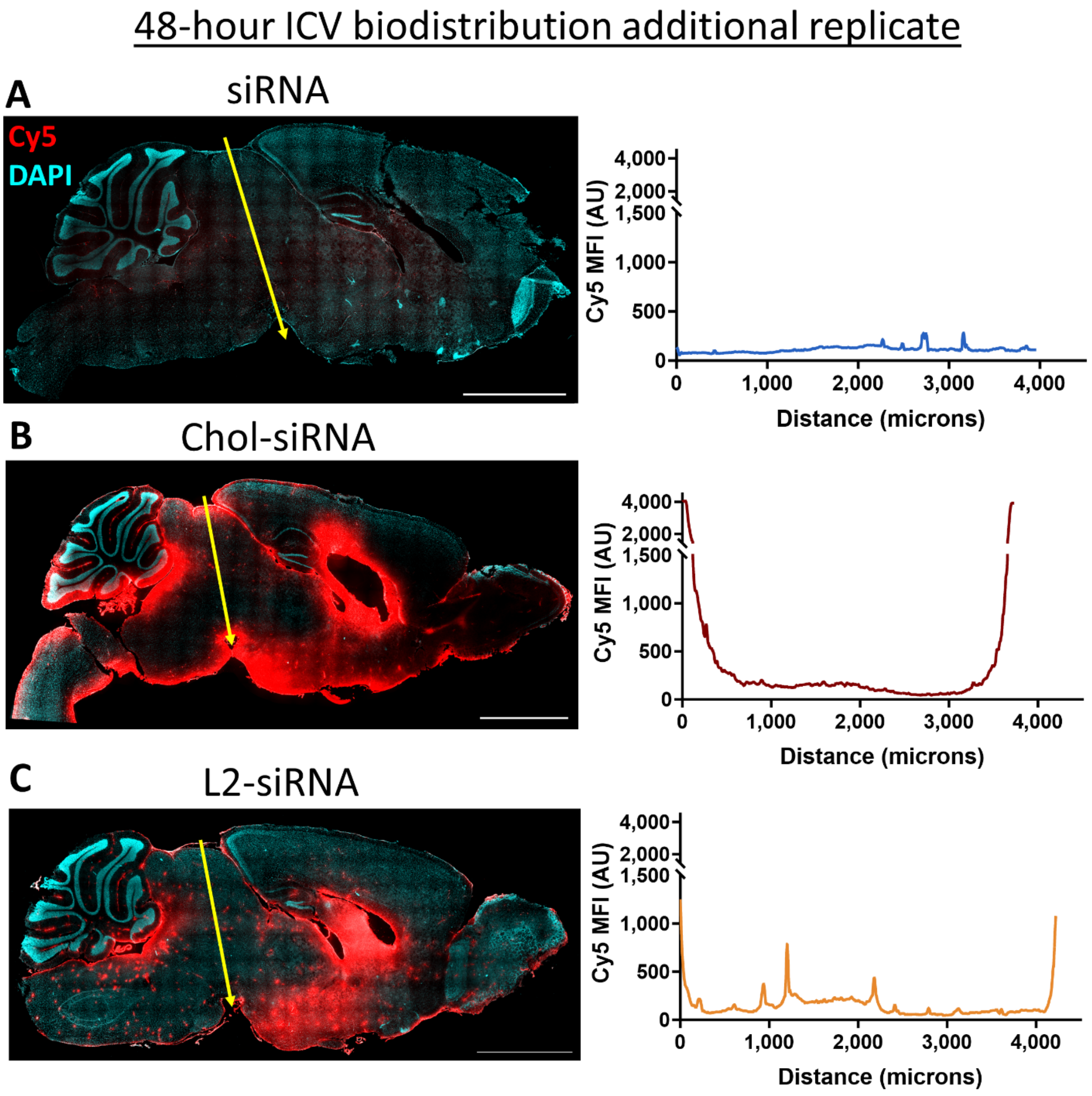

**Supplementary Figure 3: Additional replicates of biodistribution after ICV delivery**

Delivery throughout CNS 48 hours after ICV delivery. Cy5 mean fluorescence intensity (MFI) plotted along distance indicated by yellow line. Additional example shown for siRNA (A), Chol-siRNA (B), and L2-siRNA (C). Section thickness = 30 µm, scale bar = 2.5 mm.

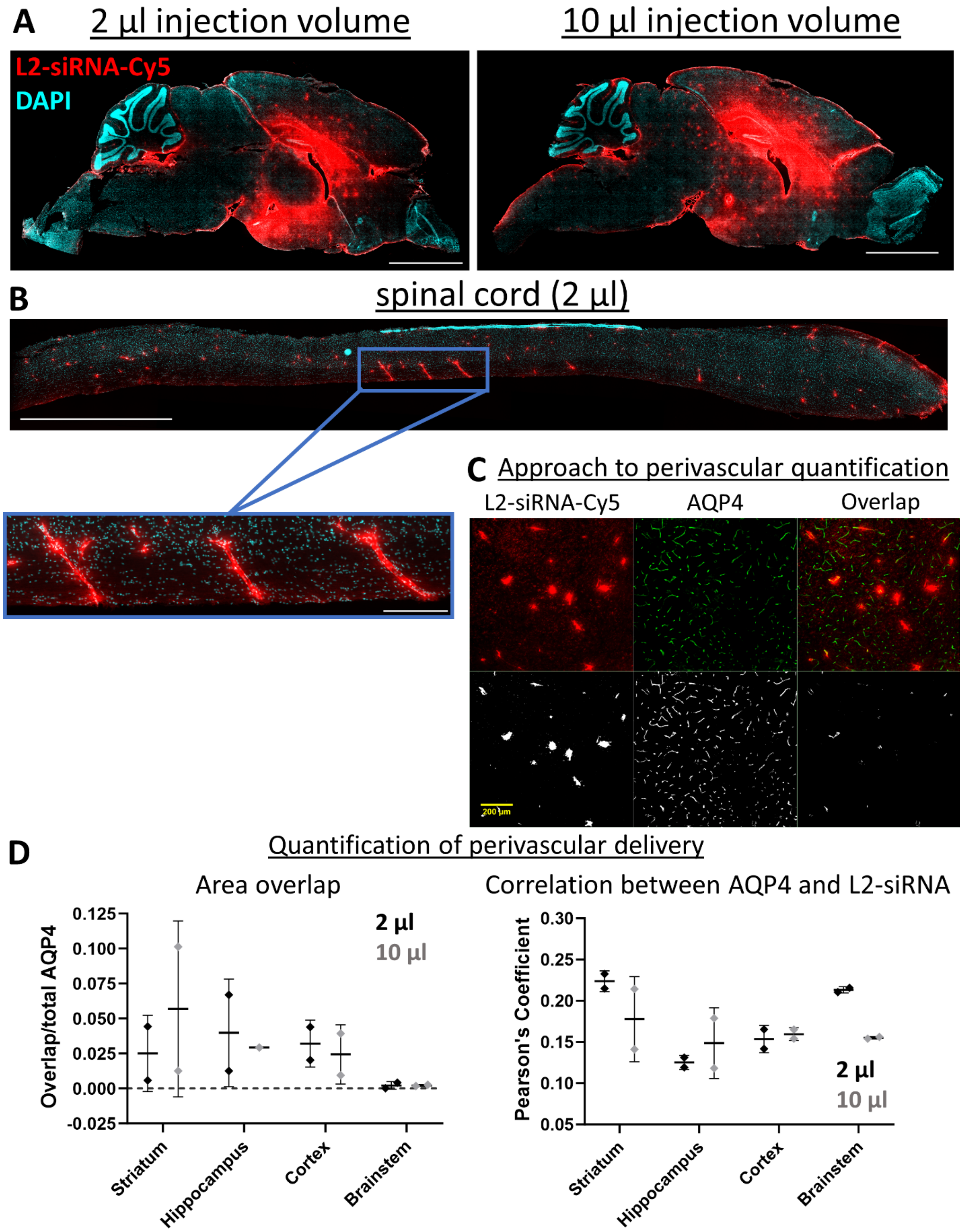

A
2 µl injection volume
10 µl injection volume
L2-siRNA-Cy5
DAPI
B
spinal cord (2 µl)
C
Approach to perivascular quantification
L2-siRNA-Cy5
AQP4
Overlap
200 µm
D
Quantification of perivascular delivery
Area overlap
Overlap/total AQP4
0.125
0.100
0.075
0.050
0.025
0.000
-0.025
2 µl
10 µl
Striatum
Hippocampus
Cortex
Brainstem
Correlation between AQP4 and L2-siRNA
Pearson's Coefficient
0.30
0.25
0.20
0.15
0.10
0.05
2 µl
10 µl
Striatum
Hippocampus
Cortex
Brainstem

**Supplementary Figure 4: L2-siRNA delivery to perivascular spaces is observed with low injection volumes**

A, B. To test whether perivascular delivery is an artifact of injection volume, mice were administered an equivalent ICV dose of L2-siRNA (2 nmol), in either 2 µl or 10 µl by adjusting the initial compound concentration appropriately. After 48 hours, biodistribution was examined in the parenchyma (A) and spinal cord (B), with L2-siRNA shown in red, counterstained with DAPI in cyan (30 µm sections, scale bar = 2.5 mm).

C, D. Quantification of L2-siRNA delivery to perivascular spaces computed as the colocalization of AQP4, demarcating the outer boundary of the PVS, with Cy5-tagged L2-siRNA. Area overlap is determined by the Mander's coefficient, which reflects the fraction of AQP4 staining that is Cy5 positive. The overall association between the signals was determined using Pearson's correlation with predefined regions of interest for different brain regions. Scale bar = 200 µm. N=2 mice per treatment, mean ± SD.

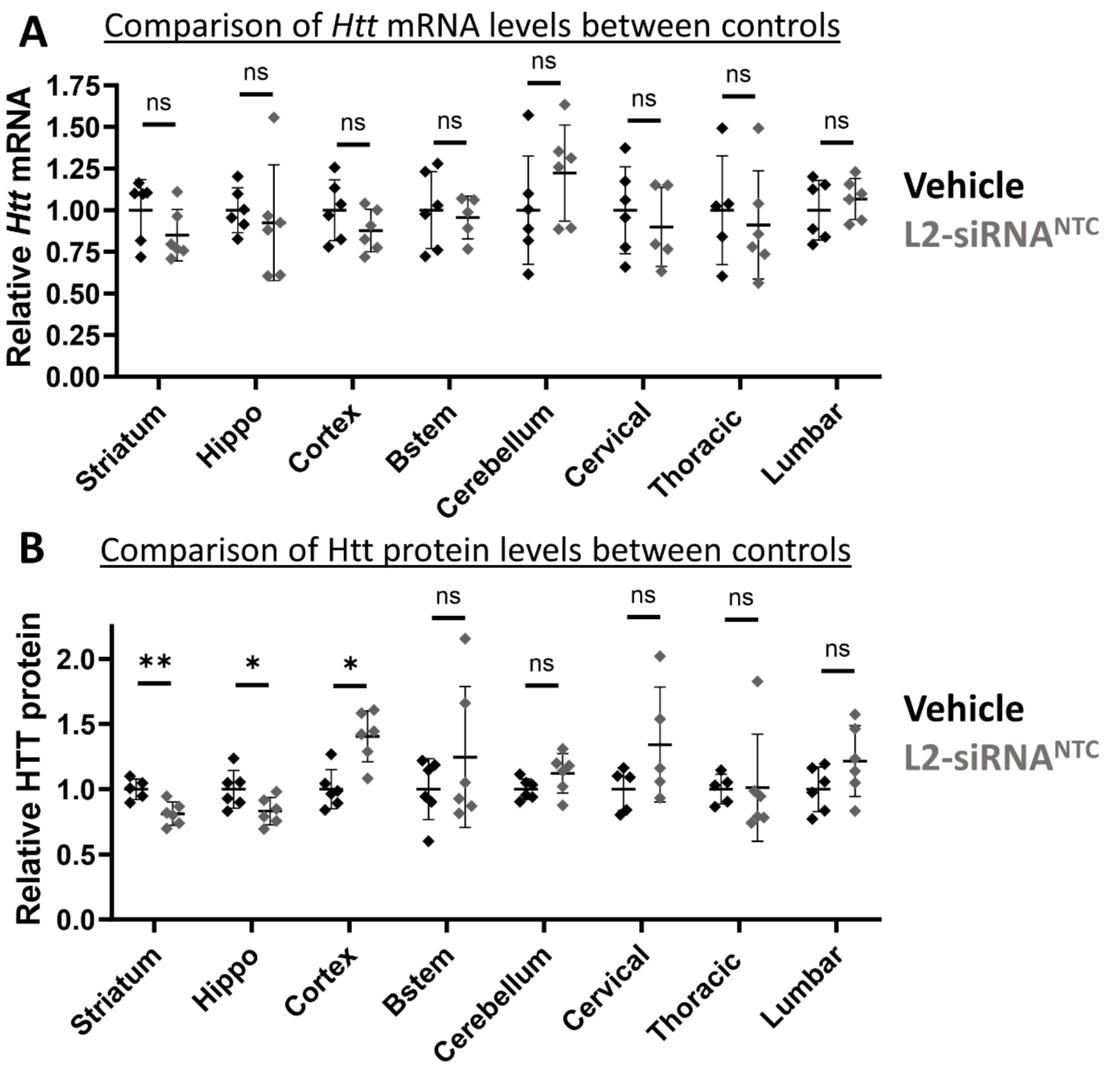

**Supplementary Figure 5: Comparison of Htt expression between negative controls**

Knockdown of Htt was assessed one month after ICV injection of vehicle (0.9% NaCl) or L2-siRNA$^{NTC}$ (15 nmol) at the mRNA level by RT-qPCR (A) and protein level by western blot (B). Each data point represents one mouse (N=6), is normalized to vehicle, and represented as mean ± SD. Statistics were computed as unpaired, two-tailed t-tests (ns – not significant, *$p<0.05$, **$p<0.01$). Hippo = hippocampus, Bstem = brainstem.

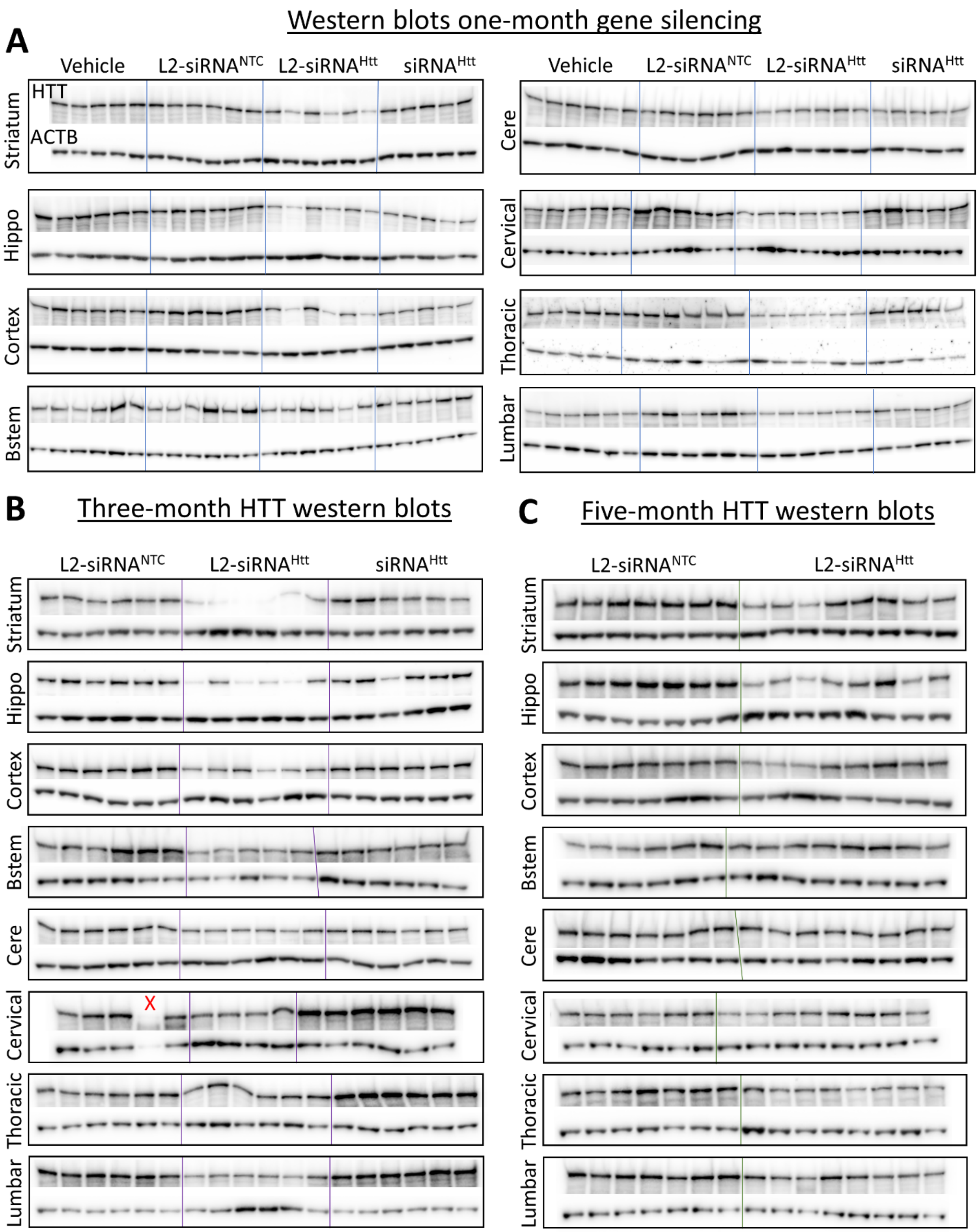

Western blots one-month gene silencing
A
Vehicle
L2-siRNANTC
L2-siRNAHtt
siRNAHtt
HTT
ACTB
Striatum
Hippo
Cortex
Bstem
Cere
Cervical
Thoracic
Lumbar
B
Three-month HTT western blots
C
Five-month HTT western blots
X

**Supplementary Figure 6: Raw western blots to assess HTT protein silencing**

Western blots associated with protein knockdown quantified in Figure 2C. For all boxes, the top bands are HTT and the bottom bands are the housekeeping gene beta-actin (ACTB). Each band represents an individual mouse (i.e. biological replicate). Blots are shown for each region and timepoint. Red X marks a sample with degraded protein that was excluded from analysis.

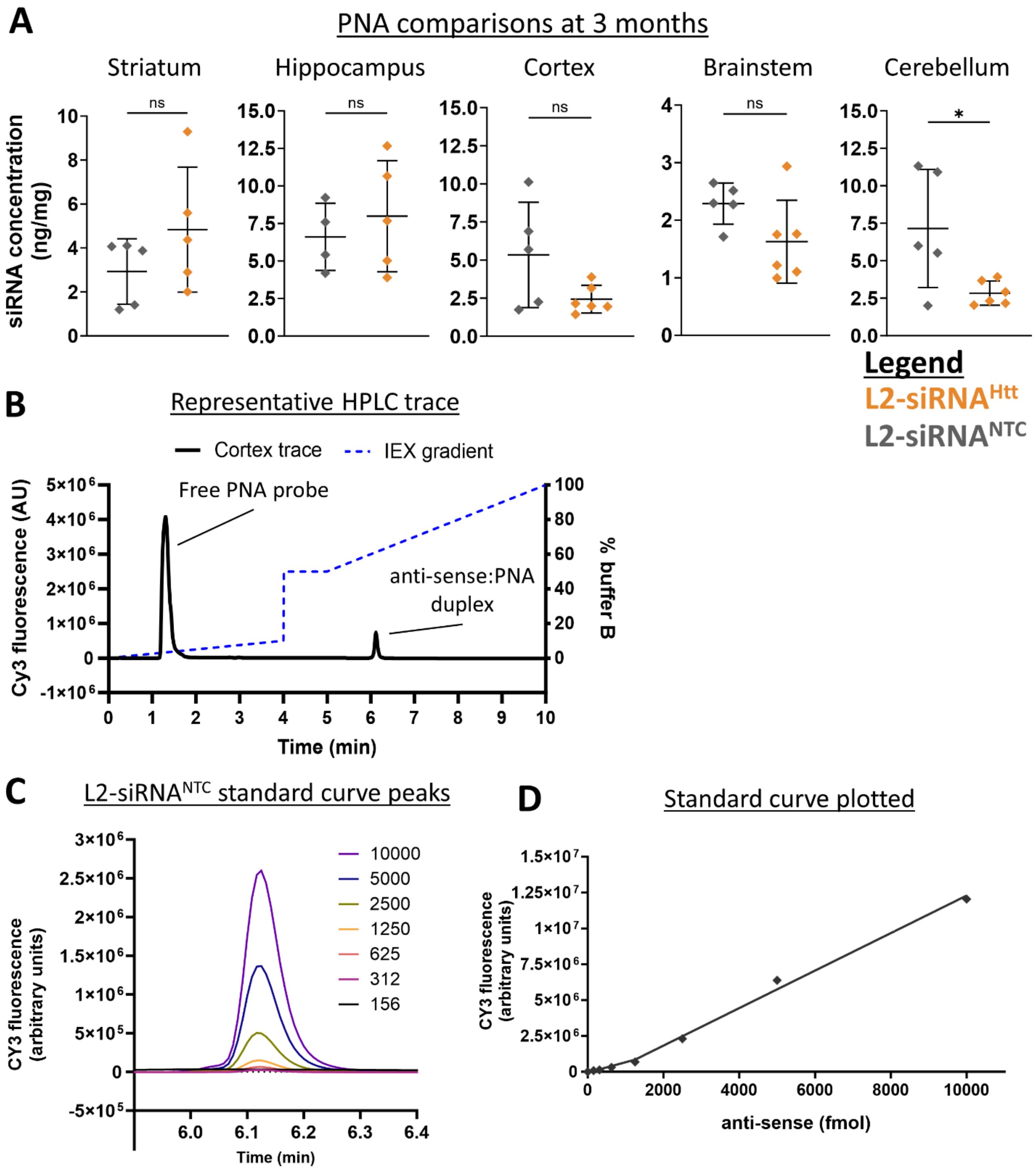

A
PNA comparisons at 3 months
Striatum
Hippocampus
Cortex
Brainstem
Cerebellum
siRNA concentration (ng/mg)
ns
ns
ns
ns
*
Legend
L2-siRNAHtt
L2-siRNANTC
B
Representative HPLC trace
Cortex trace
IEX gradient
Free PNA probe
anti-sense:PNA duplex
Cy3 fluorescence (AU)
% buffer B
Time (min)
C
L2-siRNANTC standard curve peaks
10000
5000
2500
1250
625
312
156
CY3 fluorescence (arbitrary units)
Time (min)
D
Standard curve plotted
CY3 fluorescence (arbitrary units)
anti-sense (fmol)

**Supplementary Figure 7: PNA assay supporting information**

A. Comparing siRNA delivery between L2-siRNA$^{NTC}$ and L2-siRNA$^{Htt}$ 3 months after ICV injection (15 nmol). Units are nanograms (ng) of anti-sense strand per milligram (mg) of tissue. Mean ± SD from N=5-6 biological replicates, two-tailed unpaired t-tests for each region (ns – not significant, *p<0.05).

B. Representative HPLC trace showing elution of anti-sense:PNA complex overlayed with ion-exchange gradient (dashed blue line). Trace shown from a cortex sample of L2-siRNA$^{NTC}$.

C. Raw peaks for L2-siRNA$^{NTC}$ standard curve (in fmol units).

D. Quantification of peaks represented with a bilinear fit for L2-siRNA$^{NTC}$ standard curve.

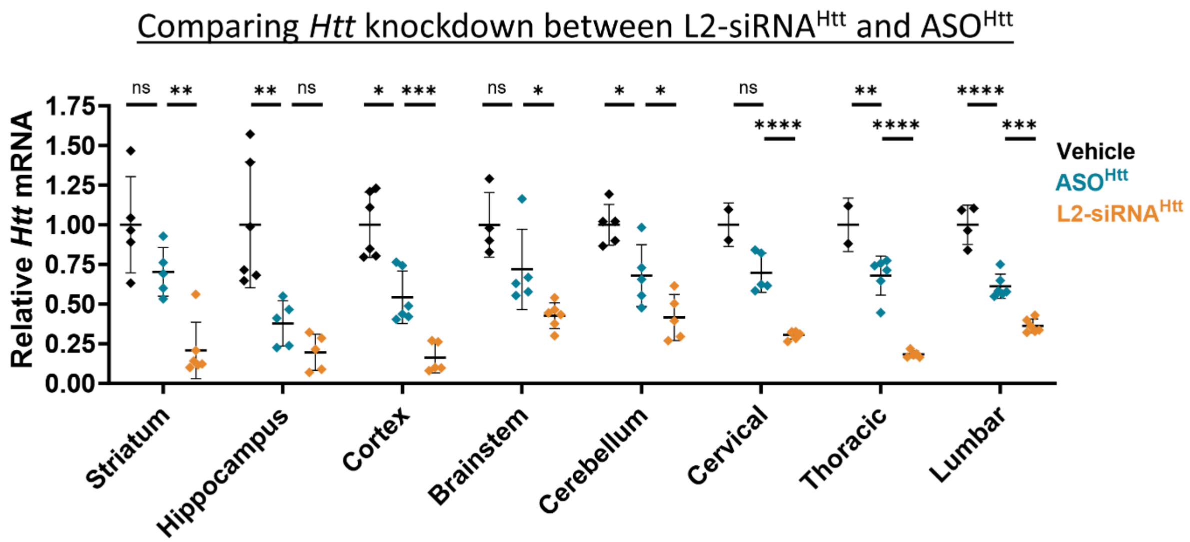

**Supplementary Figure 8: Comparing Htt-EG18 silencing of *htt* with preclinical ASO.**

Mice were administered 15 nmol of $ASO^{Htt}$ or L2-$siRNA^{Htt}$ ICV and knockdown was normalized to a vehicle (0.9% NaCl) control. The $ASO^{Htt}$ sequence targeting was designed elsewhere[1] and ordered from IDT. *Htt* levels were determined by RT-qPCR one month after injection. Since the $ASO^{Htt}$ and L2-$siRNA^{Htt}$ knockdown studies were performed on different days, each sample is normalized to their respective cohort of vehicle controls, and only the vehicle from the $ASO^{Htt}$ study is plotted here. Each point represents a mouse (N=6) and bars represent mean ± SD. One-way ANOVA was performed for each region comparing $ASO^{Htt}$ to vehicle, L2-$siRNA^{Htt}$ to $ASO^{Htt}$ (Bonferroni's correction for multiple comparison).

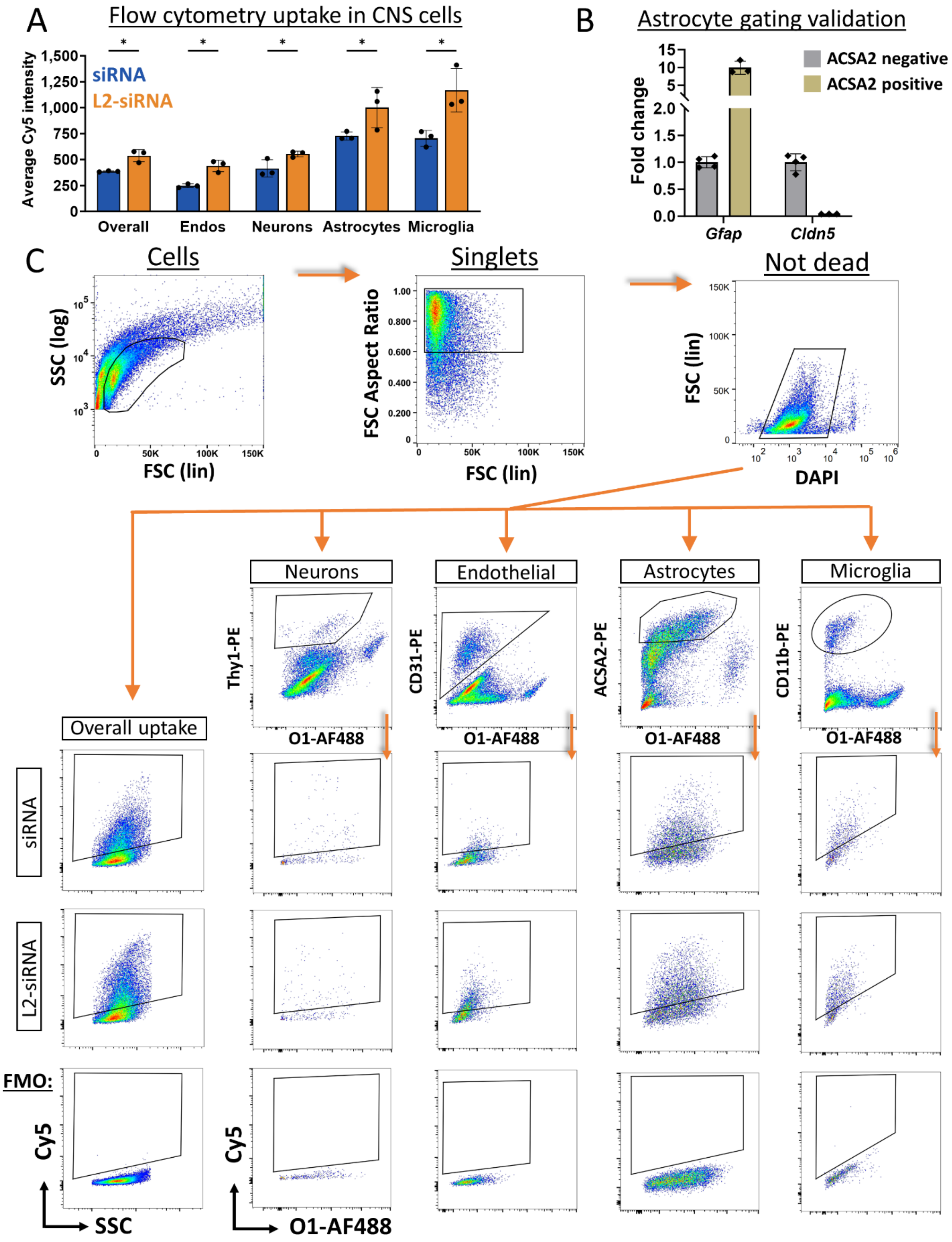
A
Flow cytometry uptake in CNS cells
Average Cy5 intensity
siRNA
L2-siRNA
Overall
Endos
Neurons
Astrocytes
Microglia
B
Astrocyte gating validation
Fold change
ACSA2 negative
ACSA2 positive
Gfap
Cldn5
C
Cells
SSC (log)
FSC (lin)
Singlets
FSC Aspect Ratio
FSC (lin)
Not dead
FSC (lin)
DAPI
Neurons
Thy1-PE
O1-AF488
Endothelial
CD31-PE
O1-AF488
Astrocytes
ACSA2-PE
O1-AF488
Microglia
CD11b-PE
O1-AF488
Overall uptake
siRNA
L2-siRNA
FMO:
Cy5
SSC
Cy5
O1-AF488

**Supplementary Figure 9: Flow cytometry assessment of cell-specific uptake**

A. Cell-specific uptake assessed 48 hours after ICV injection of L2-siRNA or siRNA (10 nmol). Data reported as mean Cy5 intensity of live cells. Statistics computed as unpaired, two-tailed t-tests corrected for multiple comparisons using the false discovery rate (FDR) approach with Q=5%. N=3 mice, data presented as mean ± SD.

B. Validation of astrocyte gating strategy by performing RT-qPCR on fluorescence activated cell sorting (FACS) purified ACSA2+/O1- and ACSA2-/O1- cells.

C. Gating strategy for identifying cell types and measuring Cy5 uptake. All samples are pre-gated to separate cells from debris, single cells from doublets, and live cells from DAPI+ dead cells. Neurons are defined as Thy1+/O1-, endothelial cells as CD31+/O1-, Astrocytes as ACSA2+/O1, Microglia/macrophages as CD11b+/O1-. Within these cell populations, Cy5+ events are gated using fluorescence-minus one (FMO) controls unique to each cell type. Representative plots are shown (median of each triplicate). AF488 = alexafluor 488, PE = phycoerythrin, FSC = forward scatter, SSC = side scatter.

**A** Quality control and filtering of CD11b$^{pos}$ cells

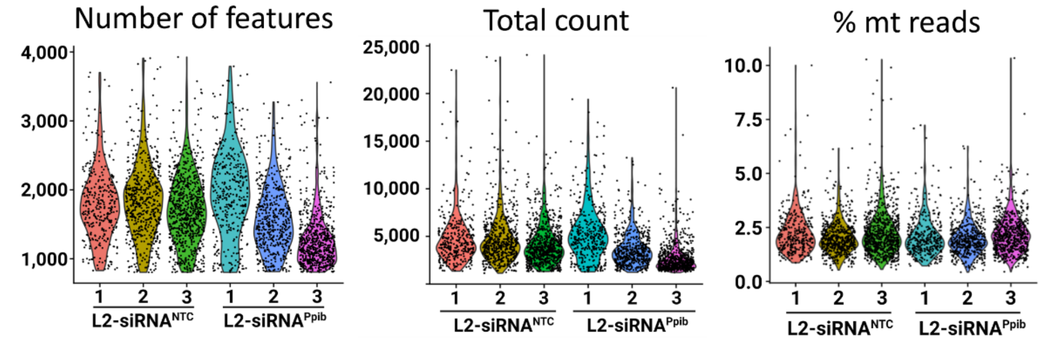

**B** Cell-type annotation

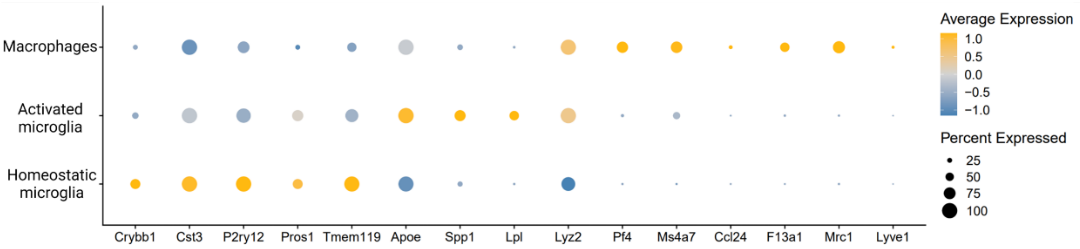

**C** Table of cell yield per sample

| | Total CD11b+ | Homeostatic microglia | Activated microglia | Macrophages | Unknown |
|---|---|---|---|---|---|
| L2-siRNA$^{NTC}$ 1 | 343 | 293 | 28 | 18 | 4 |
| L2-siRNA$^{NTC}$ 2 | 573 | 480 | 53 | 29 | 11 |
| L2-siRNA$^{NTC}$ 3 | 771 | 645 | 72 | 49 | 5 |
| L2-siRNA$^{Ppib}$ 1 | 368 | 319 | 25 | 23 | 1 |
| L2-siRNA$^{Ppib}$ 2 | 505 | 437 | 24 | 33 | 11 |
| L2-siRNA$^{Ppib}$ 3 | 701 | 615 | 25 | 51 | 10 |

**D** *Ppib* expression density plots

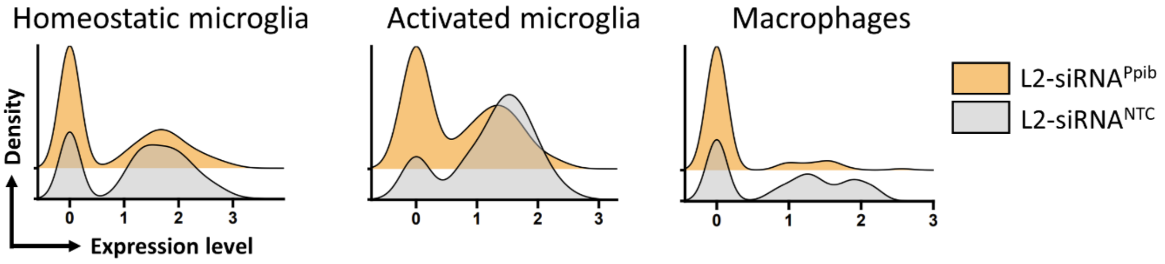

**Supplementary Figure 10: CD11b$^{pos}$ scRNA-seq supporting information**

A. Standard quality control metrics were employed for each sample processed with T2 PIPseq kits. Cells were filtered based on number of features, keeping those containing 800-4,000 uniquely expressed genes. Total count represents the absolute number of transcripts detected and percent mitochondrial (% mt) reads is the fraction of transcripts associated with mitochondrial mRNA.

B. Dotplot of canonical Cd11b$^{pos}$ population markers.

C. Number of cells identified in each CD11b$^{pos}$ population for all replicates.

D. Smoothed ridgeline density plots showing changes in *Ppib* expression between L2-siRNA$^{NTC}$ and L2-siRNA$^{Ppib}$.

**A** Quality control and filtering of CD11b$^{neg}$

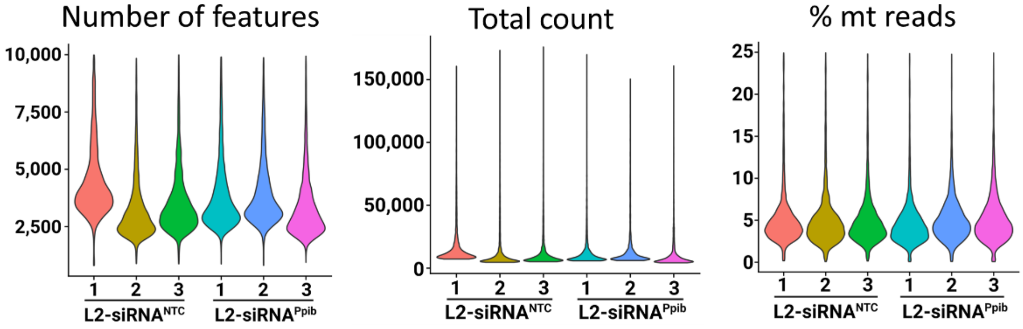

**B** Table of cell yield per sample

| | L2-siRNA$^{NTC}$ | L2-siRNA$^{Ppib}$ | | L2-siRNA$^{NTC}$ | L2-siRNA$^{Ppib}$ |
|---|---|---|---|---|---|
| Smooth muscle cells | 221 | 300 | Dural border cells | 508 | 368 |
| Endo/Pericytes | 284 | 231 | Inner arachnoid cells | 107 | 108 |
| Endo (arterial) | 446 | 517 | Arachnoid barrier cells | 33 | 35 |
| Endo (venous/capillary) | 1015 | 1310 | Parenchymal perivascular fibroblasts | 64 | 77 |
| ChP epithelial cells | 2481 | 2259 | Pial fibroblasts | 50 | 48 |
| Ependymal cells | 2725 | 2195 | Olfactory ensheathing cells | 101 | 353 |
| Oligodendrocytes | 2548 | 2368 | Neural IPCs | 359 | 308 |
| Astrocytes | 770 | 1965 | Neurons | 182 | 104 |
| Bergmann glia | 960 | 416 | Microglia | 62 | 50 |

**C** Ridgeline plots of *Ppib* expression

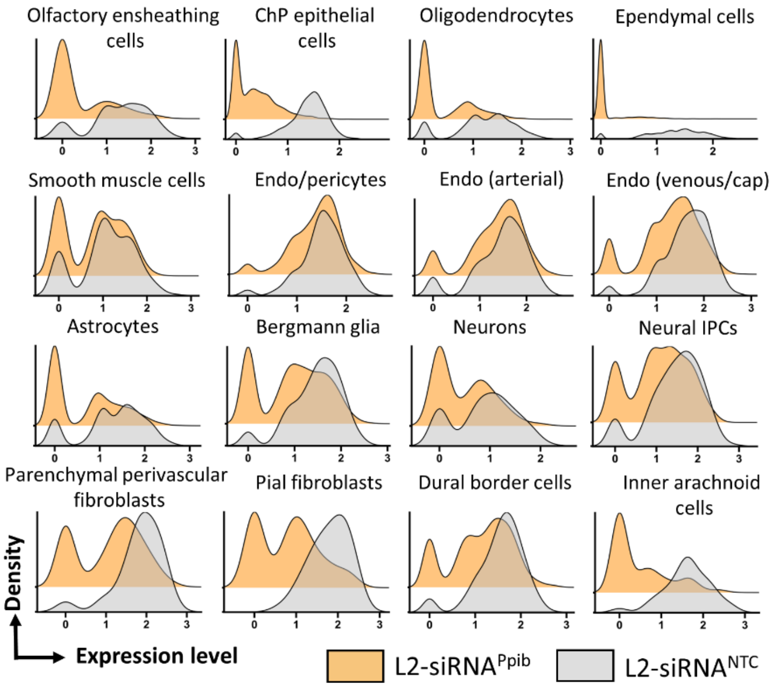

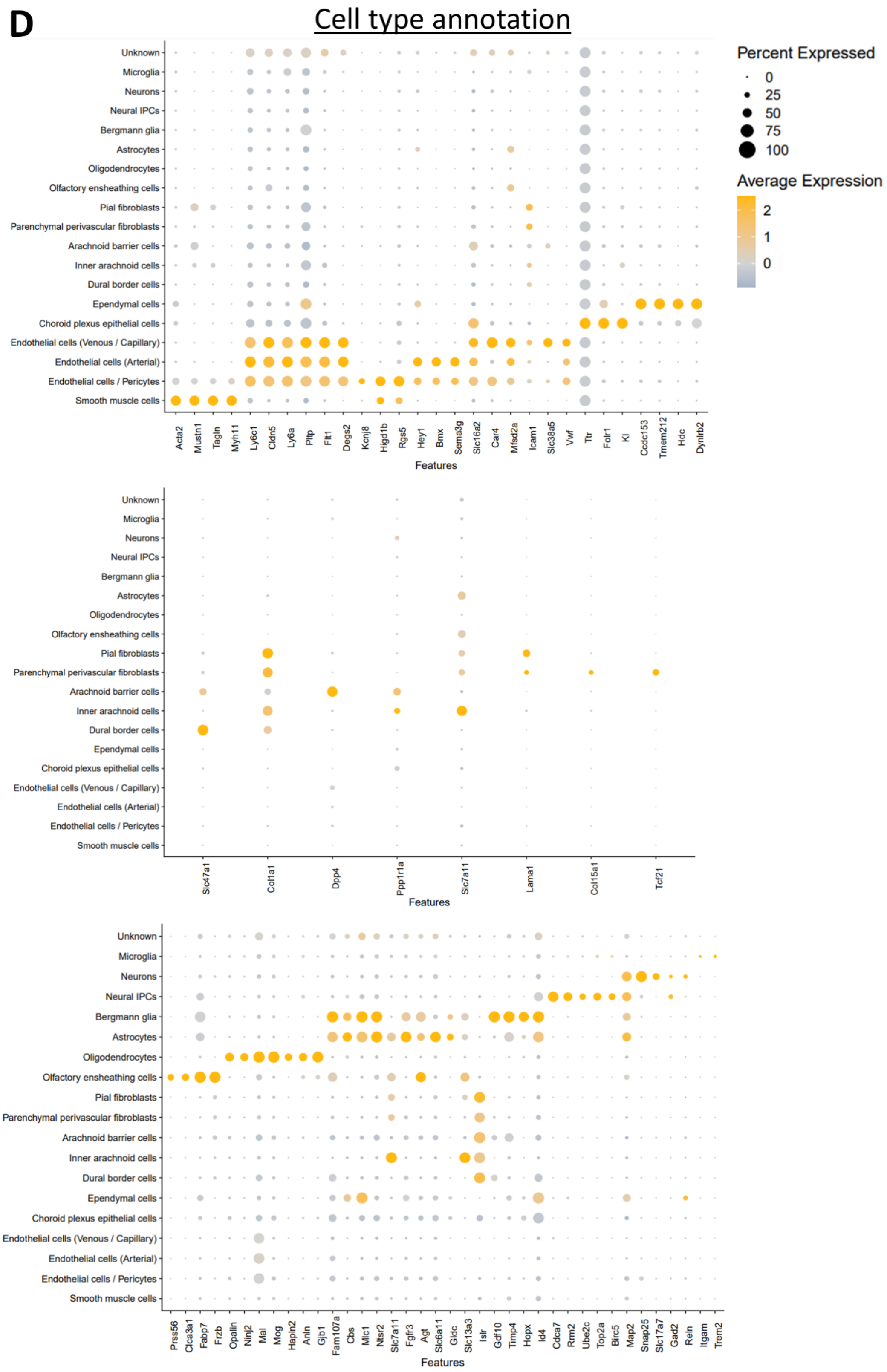

D
Cell type annotation
Percent Expressed
0
25
50
75
100
Average Expression
2
1
0
Unknown
Microglia
Neurons
Neural IPCs
Bergmann glia
Astrocytes
Oligodendrocytes
Olfactory ensheathing cells
Pial fibroblasts
Parenchymal perivascular fibroblasts
Arachnoid barrier cells
Inner arachnoid cells
Dural border cells
Ependymal cells
Choroid plexus epithelial cells
Endothelial cells (Venous / Capillary)
Endothelial cells (Arterial)
Endothelial cells / Pericytes
Smooth muscle cells
Acta2
Mustn1
Tagln
Myh11
Ly6c1
Cldn5
Ly6a
Pltp
Flt1
Degs2
Kcnj8
Higd1b
Rgs5
Hey1
Bmx
Sema3g
Slc16a2
Car4
Mfsd2a
Icam1
Slc38a5
Vwf
Ttr
Folr1
Kl
Ccdc153
Tmem212
Hdc
Dynlrb2
Features
Slc47a1
Col1a1
Dpp4
Ppp1r1a
Slc7a11
Lama1
Col15a1
Tcf21
Features
Prss56
Clca3a1
Fabp7
Frzb
Opalin
Ninj2
Mal
Mog
Hapln2
Anln
Gjb1
Fam107a
Cbs
Mlc1
Ntsr2
Slc7a11
Fgfr3
Agt
Slc6a11
Gldc
Slc13a3
Islr
Gdf10
Timp4
Hopx
Id4
Cdca7
Rrm2
Ube2c
Top2a
Birc5
Map2
Snap25
Slc17a7
Gad2
Reln
Itgam
Trem2
Features

**Supplementary Figure 11: CD11b$^{neg}$ scRNA-seq supporting information**

A. Standard quality control metrics were examined for each sample processed with T20 PIPseq kits. Cells were filtered based on number of features, keeping those containing 800-10,000 uniquely expressed genes.

B. Table of cell yield for L2-siRNA$^{NTC}$ and L2-siRNA$^{Ppib}$, reflecting the sum of cells from N=3 samples for each treatment.

C. Smoothed ridgeline density plots of *Ppib* expression for L2-siRNA$^{NTC}$ and L2-siRNA$^{Ppib}$ in each CD11b$^{neg}$ population identified.

D. Dotplot of canonical Cd11b$^{neg}$ population markers.

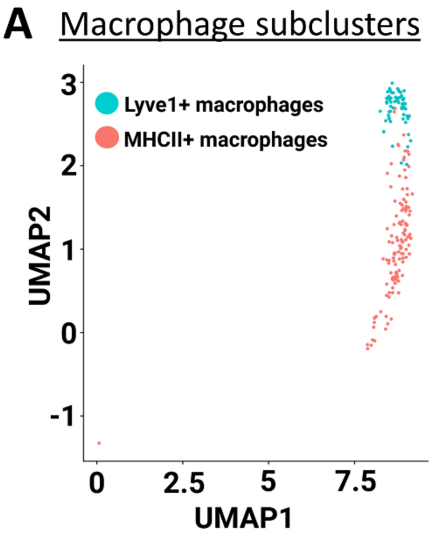

**B** Cell yield per sample

| | MHCII+ | Lyve1+ |
|---|---|---|
| L2-siRNA$^{NTC}$ 1 | 9 | 6 |
| L2-siRNA$^{NTC}$ 2 | 16 | 11 |
| L2-siRNA$^{NTC}$ 3 | 36 | 13 |
| L2-siRNA$^{Ppib}$ 1 | 13 | 8 |
| L2-siRNA$^{Ppib}$ 2 | 18 | 9 |
| L2-siRNA$^{Ppib}$ 3 | 31 | 15 |

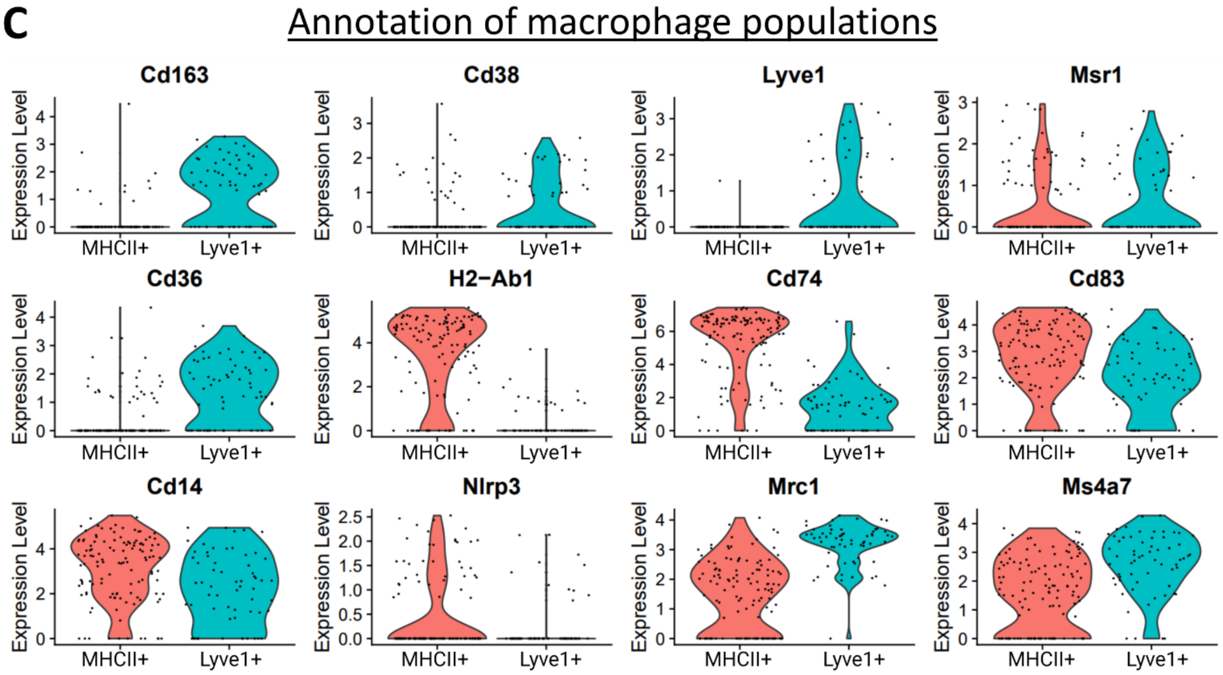

**Supplementary Figure 12: Identifying macrophage subtypes**

A. UMAP projection plot showing re-clustering of macrophages into Lyve1+ and MHCII+, as described elsewhere.[2]

B. Number of cells identified in each macrophage population for all replicates combined.

C. Annotation of gene expression in Lyve1+ and MHCII+ macrophages.

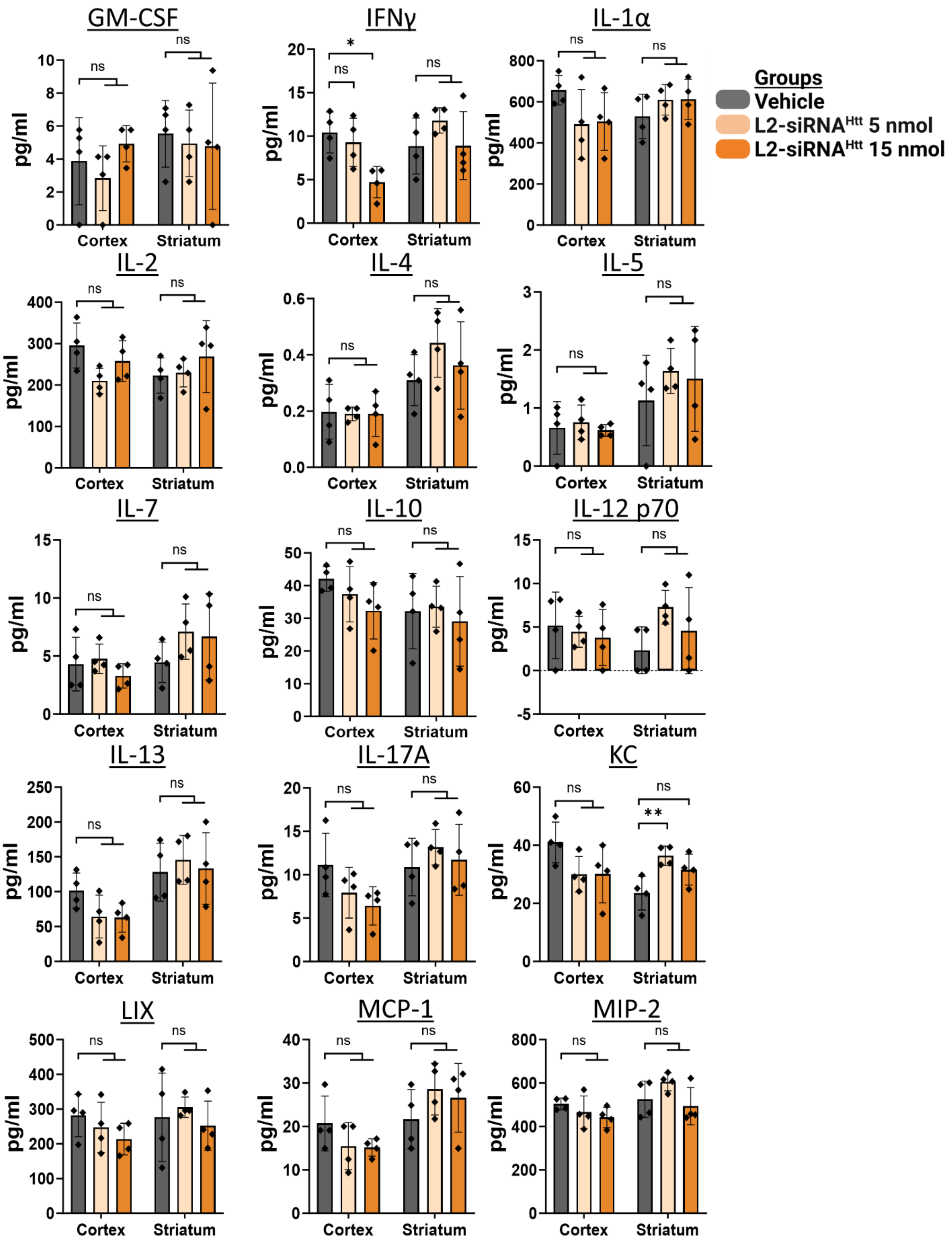
GM-CSF
IFNγ
IL-1α
Groups
Vehicle
L2-siRNAHtt 5 nmol
L2-siRNAHtt 15 nmol
IL-2
IL-4
IL-5
IL-7
IL-10
IL-12 p70
IL-13
IL-17A
KC
LIX
MCP-1
MIP-2
pg/ml
Cortex
Striatum
ns
*
**

**Supplementary Figure 13: Inflammatory protein ELISA panel**

Two weeks after ICV injection of vehicle (0.9% NaCl) or L2-siRNA$^{Htt}$ (5 nmol or 15 nmol), cortex and striatum homogenates were analyzed for presence of inflammatory proteins (cytokines, chemokines, growth factors). All samples below the limit of detection on the standard curve were treated as 0 in terms of graphical presentation and statistical analysis. All samples were below the limit of detection for TNFα, and therefore data are not shown. All data are presented as mean ± SD. A one-way ANOVA with Bonferroni's correction was performed for each region and protein (ns – not significant, *$p<0.05$, **$p<0.01$). GM-CSF, granulocyte-macrophage colony stimulating factor; IFNγ, interferon gamma; KC, keratinocyte chemoattractant; LIX, Lipopolysaccharide-induced CXC chemokine; MCP-1, monocyte chemoattractant protein-1; MIP-2, macrophage inflammatory protein-2.

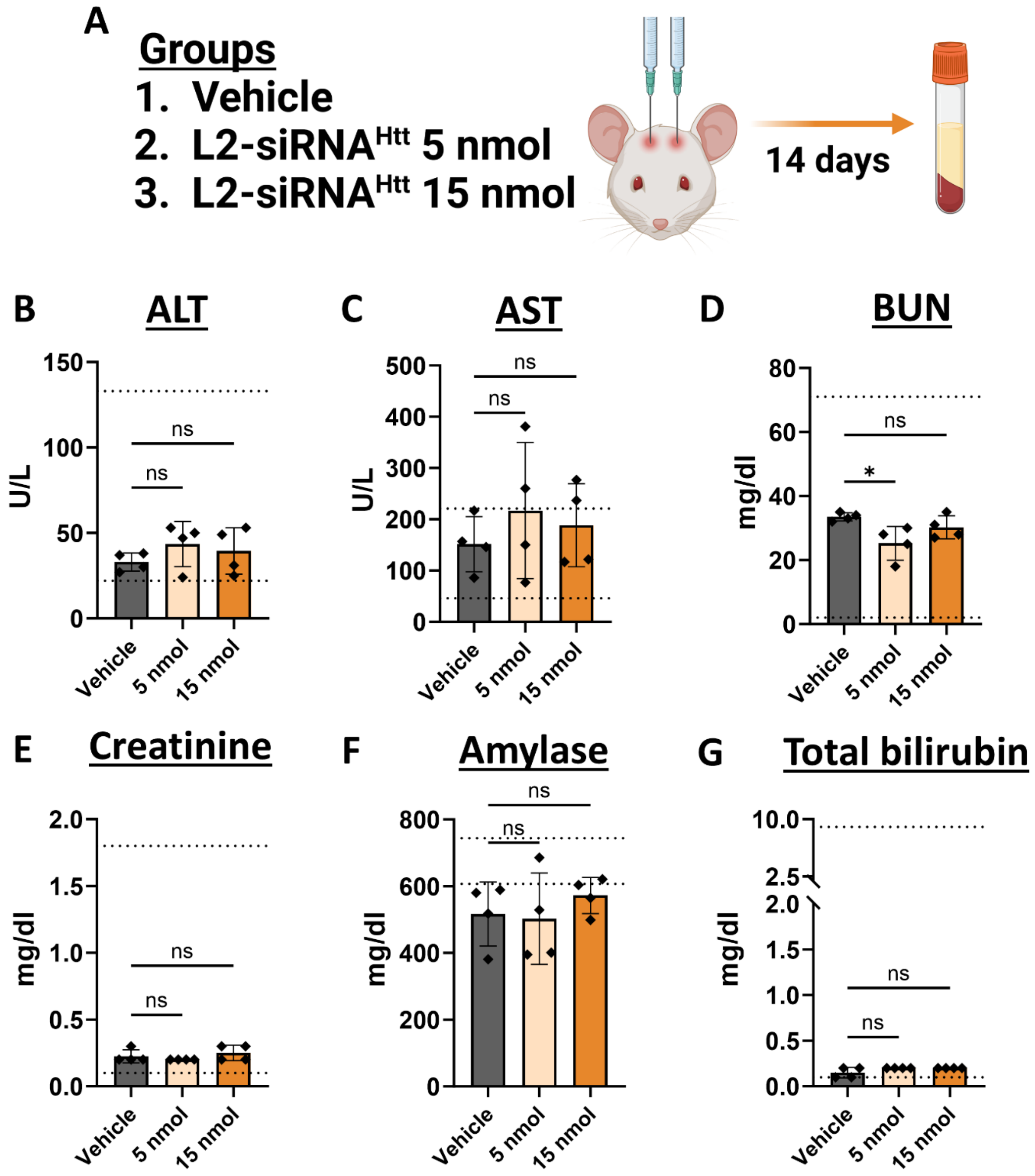

**Supplementary Figure 14: Serum chemistry assessment of organ toxicity**

Mice were injected ICV with a vehicle (0.9% NaCl) or L2-siRNA$^{Htt}$ (5 nmol or 15 nmol), and blood serum was isolated after two weeks. Six standard markers were measured in this panel: alanine aminotransferase (ALT), aspartate aminotransferase (AST), blood urea nitrogen (BUN), creatinine, amylase, and total bilirubin. Data are presented as mean ± SD. Statistics are computed as a one-way ANOVA with Bonferroni's correction compared to vehicle (ns – not significant, *$p<0.05$). Standard chemistry reference range was obtained from UCLA Division of Laboratory Animal Medicine and is represented by dotted lines.

| Name | Sequence (5’->3’) | Location | Length | Ref |
|---|---|---|---|---|
| Htt S | (MeU)*(fA)*(MeU)(fA)(MeU)(fC)(MeA)(fG)(MeU)(fA)(MeA)(fA)(MeG)(fA)(MeG)(fA)(MeU)(fU)*(MeA)*(fA) | 10150 | 20 | [24] |
| Htt AS | VP(meU)*(fU)*(MeA)(fA)(MeU)(fC)(MeU)(fC)(MeU)(fU)(MeU)(fA)(MeC)(fU)(MeG)(fA)(MeU)(fA)*(MeU)*(fA) | 10150 | 20 | [24] |
| PPIB S | (MeA)*(fA)*(MeC)(fA)(MeG)(fC)(MeA)(fA)(MeA)(fU)(MeU)(fC)(MeC)(fA)(MeU)(fC)(MeG)(fU)*(MeG)*(fA) | 437 | 21 | [34] |
| PPIB AS | VP(meU)*(fC)*(meA)(fC)(meG)(fA)(meU)(fG)(meG)(fA)(meA)(fU)(meU)(fU)(meG)(fC)(meU)(fG)*(meU)*(fU) | 437 | 21 | [34] |
| NTC S | (fC)*(MeA)*(fA) (MeU) (fU) (MeG) (fC) (MeA) (fC) (MeU) (fG) (MeA) (fU) (MeA) (fA) (MeU) (fG)*(MeA)*(fA) | NA | 19 | [29] |
| NTC AS | VP(MeU)*(fU)*(MeC) (fA) (MeU) (fU) (MeA) (fU) (MeC) (fA) (MeG) (fU) (MeG) (fC) (MeA) (fA) (MeU)*(fU)*(MeG) | NA | 19 | [29] |
| $ASO^{Htt}$ | CTCGActaaagcaggATTTC | 4042 | 20 | [8] |
| Htt PNA probe | 5’/N Cy3-OO-TATATCAGTAAAGAGATTAA 3'/C | NA | 20 | [24] |
| NTC PNA probe | 5'/N Cy3-OO-GGGACTGGCTAGTTAAACA 3'/C | NA | 19 | NA |
| Key: S=sense strand, AS=anti-sense strand, *=phosphorothioate, Me=2’-O-Methyl, f=2’fluoro, VP=vinyl phosphonate, O = ethylene glycol linker, lowercase = DNA, for $ASO^{Htt}$ uppercase = 2’-O-(2-methoxy)ethyl | | | | |

**Supplementary Table 1: Nucleic acid sequences used**
